# The impacts of seventy years of changes in stopover habitats on the critically endangered Spoon-billed Sandpiper *Calidris pygmaea*

**DOI:** 10.64898/2026.01.27.701643

**Authors:** Takehiko Shimizu, Masayuki Senzaki, Munehiro Kitazawa, Minoru Kashiwagi, Hiroshi Tomida

## Abstract

Natural habitat loss due to land use change is a major driver of global biodiversity loss. Human-created seminatural environments can function as artificial habitats for many species, partially offsetting these negative impacts. However, it remains unclear how species respond to the short– and long-term changes in both natural and artificial habitats, particularly for long-distance migratory species at stopover sites. We investigated how the globally endangered habitat specialist species, Spoon-billed Sandpipers *Calidris pygmaea*, responded to 70 years of changes in natural wetlands, sandy beaches, and artificial wetlands across stopover habitats in the Japanese Archipelago. We compiled historical observation records of the species from multiple sources and quantified the extent of these three habitat types from 1950 to 2020. Spoon-billed Sandpiper abundance consistently declined from the 1970s to the 2010s, with a particularly sharp decrease between the 1980s and the 1990s. While more than 50% of natural wetlands and sandy beaches have also been lost since 1950, we found that sandpiper abundance was lower at sites experiencing greater cumulative natural habitat loss. By contrast, changes in artificial wetland extent were not significantly associated with abundance, despite their temporal expansion peaking in the 1970s and subsequently declining. Our findings demonstrate that historical loss of natural stopover habitats have had lasting negative effects on local sandpiper populations, and the temporary expansion of artificial wetlands failed to compensate for these effects. This underscores the critical importance of preserving the networks of natural stopover habitats to sustain migration success, particularly for habitat specialists.

## 1. Introduction

Habitat loss attributed to land use change is a major driver of global biodiversity loss, even when compared with other drivers, such as climate change and pollution (Aarif et al., 2014; Horváth et al., 2019; Jaureguiberry et al., 2022; Knapp et al., 2017). Recent research combining large-scale, long-term monitoring datasets (e.g., GBIF and Living Planet Database) with land-use products (e.g., Landsat and ESA CCI LA maps) demonstrates that land use change negatively affects population size, abundance, and species richness globally (Albaladejo_Robles et al., 2023; Le Provost et al., 2020; Montràs-Janer et al., 2024; Northrup et al., 2019; Spooner et al., 2018; Williams et al., 2022). However, the magnitude of these effects can vary among habitat types within species’ ranges (Rushing et al., 2016; Williams et al., 2022) and over the time since the beginning of habitat loss (Figueiredo et al., 2019). For instance, Daskalova et al. (2020) reported that the effects of forest loss on species can lag by up to 50 years, depending on species’ generation times. More research is therefore needed to understand how species distribution relates to short– and long-term temporal changes in multiple habitat types with different functions and across large spatial scales, to inform appropriate spatial, temporal, and thematic scales for conservation actions and policy.

Artificial habitats are human-created environments, either intentionally or unintentionally (e.g., croplands, plantations, and aquaculture ponds), whereas natural habitats form through interactions between biotic and abiotic factors. Artificial habitats can function as ecological traps, reducing the fitness of species (Hale et al., 2015; Hale and Swearer, 2016; Sievers et al., 2018) and supporting lower biodiversity than natural habitats (Newbold et al., 2015). Artificial habitats are, nevertheless, increasingly recognized as potentially offsetting natural habitat loss, especially in severely degraded ecosystems such as coastal wetlands, which are critical for many aquatic species (Aarif et al., 2025; Cheng and Ma, 2023; Rajpar et al., 2022; Jackson et al., 2019; 2020). However, little is known about how species’ habitat use responds to quantitative changes in natural and artificial habitats over short– and long-term scales. Addressing this gap is essential for improving our understanding of the ecological roles of artificial habitats relative to natural ones.

Species’ responses to spatiotemporal changes across multiple habitats may also depend on where those changes occur within their ranges (Zurell et al., 2018). This is especially true for highly mobile, long-distance migratory species that use geographically distant breeding, non-breeding, and stopover sites each year (UNEP-WCMC, 2024). Stopover habitats are critical for refueling and physiological recovery and can influence migration patterns, migration success, individual fitness, and population dynamics (Lisovski et al., 2024; Liu et al., 2022; Schmaljohann et al., 2022; Studds et al., 2017). Given that migratory species experience relatively high mortality during migration (Klaassen et al., 2014), habitat loss and reduced carrying capacity at regional stopover sites may significantly affect populations at both site-specific and global scales, including subsequent stopover or non-breeding sites (Liu et al., 2022; Mu et al., 2022). Therefore, it is important to assess how short– and long-term changes in natural and artificial habitats at stopover sites limit the abundance of migratory species, especially for those with specialized ecological niches that are more vulnerable to environmental change (Bowler et al., 2018; Habel et al., 2023; Newbold et al., 2018). Nevertheless, this question remains understudied because stopover use is relatively brief compared with breeding and non-breeding periods (Flack et al., 2022).

Here, we examined how a vulnerable migratory species with specialized habitat requirements responds to spatiotemporal changes in the distribution of natural and artificial stopover habitats. We focused on Spoon-billed Sandpipers *Calidris pygmaea,* and their stopover habitats across the Japanese Archipelago. Spoon-billed Sandpipers migrate from breeding grounds in far-east Russia to non-breeding sites in southeast Asia, stopping over in East Asia, including Japan (Dixon, 1918; Jia et al., 2025; Tomkovich et al., 2002; Zöckler et al., 2016). Because their distribution is inherently limited, Spoon-billed Sandpipers are habitat specialists and are more vulnerable to habitat loss than generalist species (Zöckler et al., 2010). The global population has declined rapidly since the 1970s (Clark et al., 2018; Green et al., 2021; Tomkovich et al., 2002), leading to the species being listed as “Critically Endangered” on the IUCN Red List (BirdLife International, 2021). Spoon-billed Sandpipers also commonly forage and rest with other shorebird species, such as Red-necked Stint *C. ruficollis* and Sanderling *C. alba*. Habitat loss may therefore intensify intra– and interspecific competition within mixed-species flocks and could contribute to density-dependent population declines. Focusing on Spoon-billed Sandpipers and Japan is well suited to our study for three reasons: 1) Japan has extensive long-term historical records of this species and its habitats, enabling robust analysis; 2) natural habitats for the species—coastal wetlands such as tidal flats—have declined sharply since 1900 (Aung et al., 2022; Lu et al., 2022; Murray et al., 2022), with partial compensation from artificial wetlands (e.g., reclaimed wetlands), providing clear context to assess the impact of natural versus artificial habitat change; and 3) studying the species can provide insight into the effects of stopover habitat loss and inform conservation of this critically endangered bird.

Because global databases provide insufficient long-term observation and habitat data for this species in Japan, we compiled historical data from multiple sources and measured the extent of three habitat types—natural wetlands, sandy beaches, and artificial wetlands—from 1950 to 2020. The long-term, broad-scale dataset allowed us to address three objectives: 1) describe spatiotemporal changes in Spoon-billed Sandpiper abundance (Fig. 1A), 2) quantify changes in the area of the three habitat types (Fig. 1B), and 3) assess quantitative relationships between abundance and spatiotemporal habitat variation (Fig. 1C). We hypothesized that abundance would be higher at stopover sites with lower habitat loss (Fig. 1C). We further expected that the effects of habitat loss would vary by habitat type, original habitat extent, and time since loss began.

**Fig. 1.**
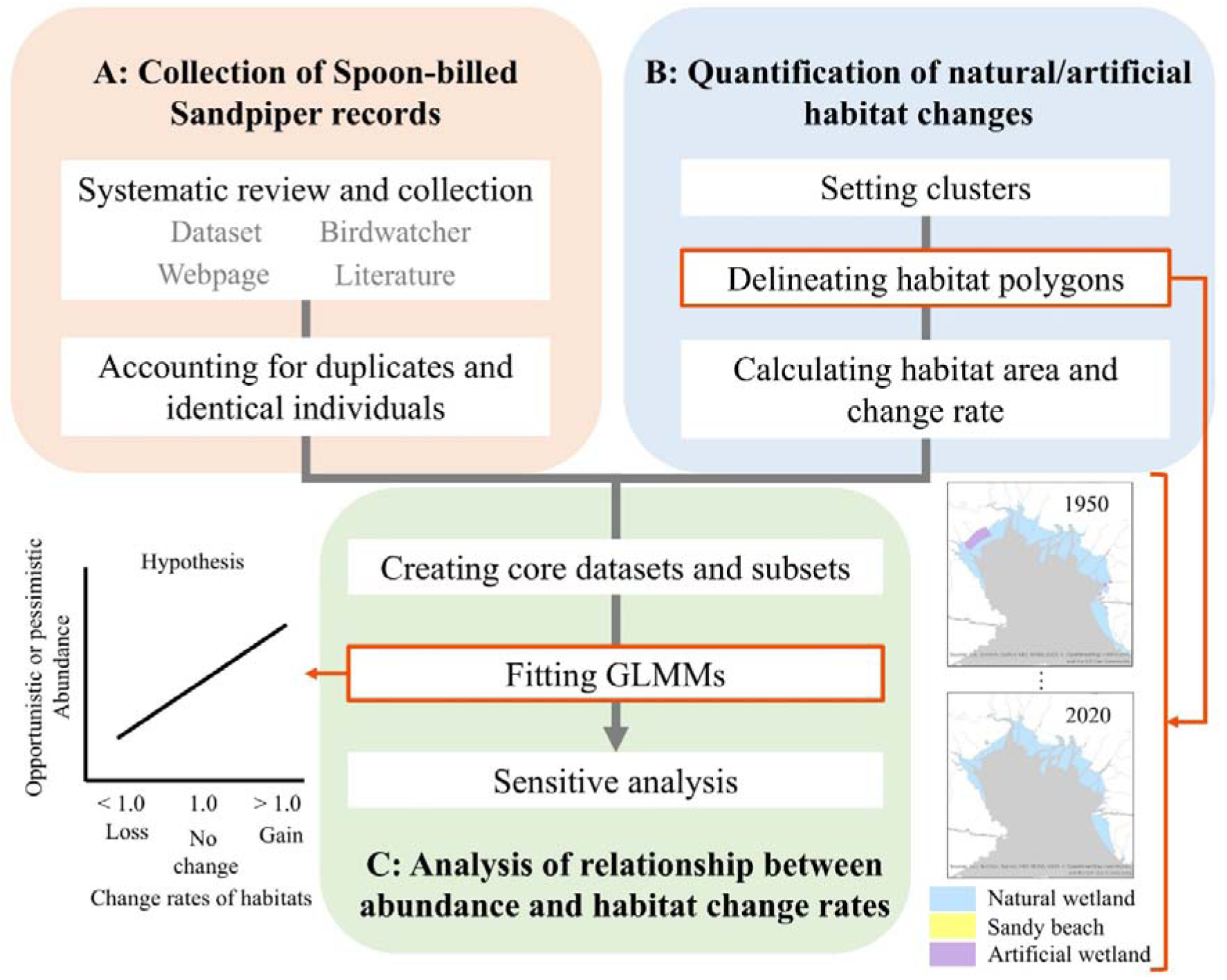
Overview of the study. A) Collection of Spoon-billed Sandpiper records to build an original database. B) Quantification of changes in three habitat types by delineating polygons at clusters where the species was observed. C) Analysis of relationships between abundance and habitat change rates.

## 2. Materials & Methods

### 2.1. Systematic collection of observation records of Spoon-billed Sandpipers

We compiled observation records of Spoon-billed Sandpipers in Japan from four main sources: public databases, local birdwatchers’ records, reliable webpages, and published literature (Table S1.1). Data compilation followed the Preferred Reporting Items for Systematic Reviews and Meta Analyses (PRISMA) guideline (Haddaway et al., 2022; Page et al., 2021) (Fig. S1.1). The three public databases were the national systematic shorebird monitoring survey *Monitoring Site 1000* (MOEJ, 2023), the Japan Banding Bird Database (https://www.birdbanding-assn.jp/), and eBird (https://ebird.org/explore). For records from local birdwatchers and webpages, we included only observations from people proficient in bird identification, verified through direct communication. Literature sources included any publication type—academic articles, newspapers, and reports—that contained observation records of Spoon-billed Sandpipers. From 24 May to 7 July 2023, we systematically collected papers using the following search platforms: J-STAGE (https://www.jstage.jst.go.jp/), Google Books (https://books.google.com/), the book search engine of the Japan Publishing Organization (https://www.books.or.jp/), and the National Diet Library, Japan (https://www.ndl.go.jp/en/index.html). We used the following Japanese keywords (shown as “English translation” [Japanese]): “Spoon-billed Sandpiper” [herashigi], “shorebird” [shigi-chidori], “bird guide book” [tori-zukan], and the names of late Japanese ornithologists listed on the Wikipedia page (“Ornithologists in Japan” [nihon-no-chōruigakusha], https://shorturl.at/1g1ht), to retrieve historical records compiled by these individuals. The complete list of keywords is provided in the Supplementary material (Table S1.1).

Additionally, we reviewed four further categories of literature to increase record coverage. The first category comprised five ornithological journals published in Japan: *Japanese Journal of Ornithology*, *Ornithological Science*, *Journal of the Yamashina Institute for Ornithology*, *Bird Research*, and *Strix*. The second category was the Red Data Book published by each prefecture in Japan. The third category was academic papers published by natural history museums in Japan, as listed by the Japanese Association of Museums (https://www.j-muse.or.jp/en/). The final category comprised bird lists compiled for each prefecture in Japan. We also included additional papers identified through reference lists during these surveys. In total, we identified 4,569 papers through the systematic survey. After screening titles, abstracts, and full texts, we retained 68 papers and added another 174 papers from other sources, resulting in 242 papers (Fig. S1.1). When integrating records from different sources, we extracted prefecture, municipality, site name, utilized habitat type (natural wetland, sandy beach, artificial wetland, inland wetland, or others; details in Section 2.2), year, month, day, number of individuals observed, and age. Missing information was left blank.

The use of the raw observation database is unsuitable for the following analysis because it contains obvious or potential duplicates. Furthermore, since the number of Spoon-billed Sandpipers per site is generally low in Japan, we used the maximum number of individuals recorded during a single stopover visit, while accounting for obviously or potentially identical individuals. Although obvious duplicates and clearly identical individuals (e.g., the same date and location, or a single individual observed on five consecutive days) could be merged into a single entry, missing information in some records prevented us from confidently identifying duplicates or identical individuals. To address this uncertainty, we summarized potentially duplicate records or identical individuals using two values: the lowest and highest plausible maximum counts. The lowest maximum value represents a “pessimistic” case, in which all potentially duplicate records or identical individuals were treated as obvious duplicates and identical individuals. On the other hand, the highest maximum value represents an “opportunistic” case, in which all potentially duplicate records or identical individuals were treated as unique records and individuals (see Supplementary materials S2 for details).

### 2.2. Quantification of natural/artificial habitat changes

We quantified changes in the area of three habitat types frequently used by Spoon-billed Sandpipers: natural wetlands, sandy beaches, and artificial wetlands (Fig. S3.1). Natural wetlands are the species’ primary foraging habitats and include tidal flats, mudflats, and estuaries. In the opportunistic and pessimistic cases, 44.0% and 45.1% of observations were made in these habitats, respectively (Table S3.1). Sandy beaches are dry beach zones primarily used as roosting sites, accounting for 22.6% and 22.9% of observations in the opportunistic and pessimistic cases, respectively (Table S3.1). Since the foreshore of sandy beaches is mainly used for foraging, we classified it as natural wetlands. We defined coastal reclaimed wetlands as artificial wetlands. Japan has historically reclaimed coastal zones for agriculture and commercial land use. During reclamation, shallow wetlands resembling natural mudflats appeared temporarily before gradually drying and becoming covered with grass or concrete. Spoon-billed Sandpipers frequently utilized these reclaimed wetlands, which accounted for 17.1% and 17.8% of observations in the opportunistic and pessimistic cases, respectively (Table S3.1). Some individuals were also observed in other types of wetlands, such as inland rice paddies. However, records in these inland wetlands were relatively infrequent (3.3–3.4%), and their spatial extent and temporal change are difficult to delineate separately from other croplands (e.g., vegetable and wheat fields) in monochrome aerial photographs; therefore, we excluded these inland records from further analysis and did not calculate the areal changes in these inland wetlands.

We defined stopover sites as contiguous areas that represent the movement ranges of Spoon-billed Sandpipers, hereafter referred to as clusters. We also defined stopover habitats as polygons representing the three types of available habitats within clusters, delineated visually and manually using satellite images and aerial photographs. We initially created clusters as 10-km-radius circles centered on each observation location, based on approximate shorebird home ranges (Choi et al., 2014). Although the cluster size represents potential movement ranges, they may vary among regions (Choi et al., 2014). Movement ranges may be limited in isolated small wetlands but may cover several tens of square kilometers in large mudflats and extensive sandy beaches. For example, one individual in our dataset moved among multiple habitats up to 30 km apart in Tokyo Bay (Wild Bird Society of Japan Tokyo, 2003). To account for regional variations in movement ranges, we merged overlapping cluster circles into a single cluster to reflect habitat continuity across broader habitat areas.

We delineated habitat polygons within each cluster for eight representative years with available reference images: 1950, 1960, 1970, 1980, 1990, 2000, 2010, and 2020. The time range captures coastal environmental change from immediately after World War II to recent years, including development associated with Japan’s rapid economic growth in the 1950s–1970s (Ito and Hoshi, 2020). We excluded inland clusters without coastlines and several small island clusters lacking aerial photographs, resulting in 80 clusters in total (Fig. S6.1). When reference images for a representative year were unavailable, we used the closest available images. After delineating polygons, we calculated the area of each habitat type within each cluster for each representative year. Polygons were created in ArcGIS Pro (Esri, 2024) using default ArcGIS satellite imagery, Google Earth Pro (https://support.google.com/earth/answer/21955?hl=en), and the Aerial Photographs Service of the Geospatial Information Authority of Japan (GIAJ, 2024) as reference sources.

### 2.3. Statistical analysis

We examined spatiotemporal observation trends of Spoon-billed Sandpipers using abundance per visit per cluster, defined as the maximum number of individuals observed simultaneously at a stopover cluster during the period from arrival to departure. We represented abundance in two ways: pessimistic (lowest) and opportunistic (highest) abundance, accounting for obviously or potentially identical individuals (Supplementary materials S2). We first described decadal trends in the total number of observation records, observed clusters, and pessimistic/opportunistic individuals across Japan (Fig. S6.2). We also summarized spatiotemporal trends in mean abundance per visit for each decade and cluster.

We integrated the species and habitat datasets and fitted generalized linear mixed models (GLMMs) to test whether spatiotemporal changes in habitat area were associated with the abundance of Spoon-billed Sandpipers (Northrup et al., 2019). Pessimistic and opportunistic abundance per visit were used as the response variable, assuming a negative binomial distribution to address overdispersion (Hartig, 2022). Fixed effects included five groups of variables: habitat area, change rates from baseline years, observation year, latitude, and survey effort. We included the log-transformed habitat area in 1950 and during the observation period. For each habitat type, change rates were calculated as the log-transformed ratio of habitat area in the observed decade to that in three baseline decades: 1950, and one and two decades prior to the observed decade. To avoid division by zero, we added 0.01 ha to all habitat areas before calculation. When identical individuals were observed in more than two clusters, we used the average habitat area and change rate across those clusters. Since the global population size estimated by previous studies was strongly correlated with year (r = −0.83) (Fig. S4.1), we included year as a fixed effect to distinguish habitat change effects from global population trends. Latitude at observation locations was included to account for regional differences in observations not explained by habitat extent or change. Survey effort was represented as the number of observations per year within each cluster to account for differences in survey intensity among years and clusters. We also included two interaction types: 1) habitat area × change rate, and 2) year × change rate, to test whether the effects of habitat changes varied with habitat extent or over time.

We checked variance inflation factors (VIFs) and correlation coefficients (r) among explanatory variables to reduce multicollinearity, excluding variables with VIF > 2 or r > 0.6 (O’brien, 2007; Zuur et al., 2010). Final fixed effects included year, latitude, survey effort, habitat area in observation for natural wetlands, sandy beaches, and artificial wetlands; change rates from 1950 for natural wetlands and sandy beaches; change rates from one decade earlier for total habitats, sandy beaches, and artificial wetlands; year × change rate interactions for four habitat types; and habitat area × change rate interactions for natural wetlands, sandy beaches, and artificial wetlands. All fixed effects were standardized before parameter estimation, except for the year which was centered. Location ID and cluster ID were included as random intercepts to deal with pseudo-replication and unmeasured heterogeneity among observation locations and clusters. As an offset, we included the scaled log-transformed cluster area to account for the difference in observation range among clusters.

Although the model accounted for spatiotemporal differences in observation frequency, our presence-only dataset could not fully capture variation in survey efforts (including both detections and non-detections) and may therefore bias results. Nevertheless, Japan has conducted regular nationwide shorebird monitoring surveys since the 1970s, including the Monitoring Site 1000 program (MOEJ, 2023), and 62% of our clusters include at least one monitoring site. Our compiled dataset also showed a relatively stable number of records between the 1970s and 2010s (Fig. S6.2A). We therefore defined a core dataset with relatively comparable survey efforts among clusters and years and conducted a sensitivity analysis by evaluating the consistency of model results between the core dataset and other subsets (see Supplementary materials S5 for details). The core dataset focused on clusters containing monitoring sites, autumn observations, and the period from the 1970s to the 2010s.

Sensitivity analyses demonstrated that two core-dataset models, using pessimistic and opportunistic abundance, produced generally consistent effect directions and magnitudes compared with models fitted to other subsets, while *p*-values differed slightly among models, particularly in terms of types of abundance indices and clusters (see Supplementary materials S7 for details). Given the robustness of results across models, we only present results from these two core-dataset models.

After fitting the models, we assessed spatial autocorrelation using Moran’s I and checked for overdispersion (Hartig, 2022). None of the models showed overdispersion, and spatial autocorrelation was weak or non-significant in all models (absolute values < 0.19) (Table S5.2). We also examined the model fit using marginal and conditional pseudo-R^2^ (Table S5.2). All analyses were conducted in R v4.4.1 (R Core Team, 2024), using the package “glmmTMB” for GLMMs (Brooks et al., 2017), “emmeans” to estimate marginal means of fixed effects (Lenth, 2024), and “DHARMa” to assess overdispersion and spatial autocorrelation (Hartig, 2022).

## 3. Results

After accounting for potentially duplicate records and identical individuals, we obtained 1,462–1,534 observation records, representing 1,194–1,418 individuals recorded from 1874 to 2024 across 257 locations in Japan (Fig. S6.1). The number of observation records—including repeated observations of the same individuals across multiple days—increased rapidly in the 1970s, possibly reflecting the start of regular monitoring surveys and improved data sharing (Fig. S6.2A). However, observation records decreased in the 2010s, potentially reflecting population declines in Japan and globally (Fig. S6.2A). Between the 1970s and 2010s, when record numbers were broadly comparable, the total number of observed clusters, total individuals, and average abundance per visit decreased (Fig. S6.2BC), with the steepest declines beginning in the 1980s (Fig. 2). For instance, average abundance per visit exceeded two in 14–20 of 44 clusters during the 1980s, but only in 4–9 of 35 clusters during the 1990s (Fig. 2). These results indicate a consistent nationwide decline in the number of Spoon-billed Sandpipers.

**Fig. 2.**
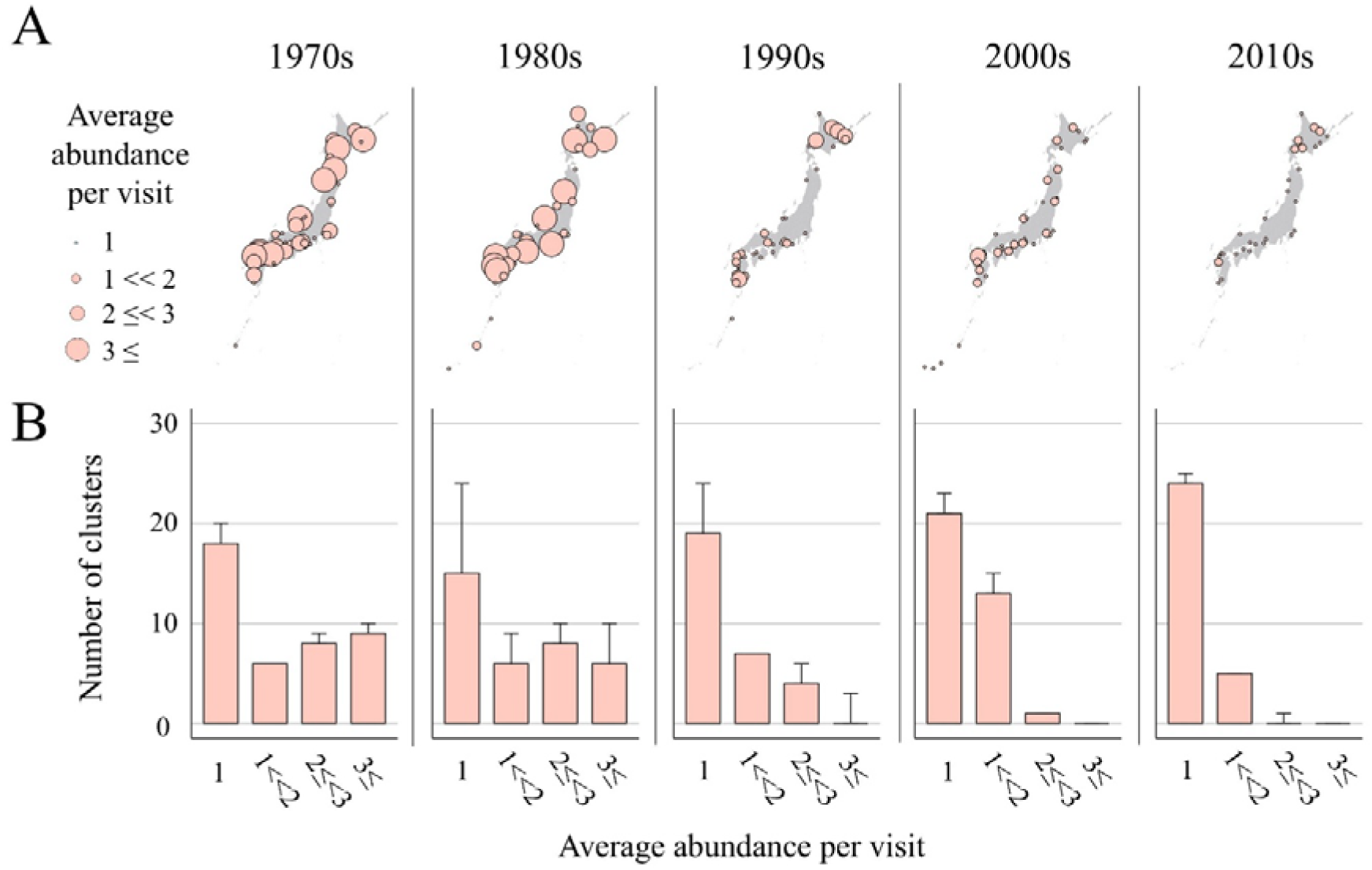
A) Maps showing the spatial distribution of average abundance per visit across five decades. B) Bar charts showing the distribution of clusters by average abundance for each decade. Bar tops denote minimum counts; error-bar tops denote the maximum count, accounting for potential identical individuals.

Habitat areas suitable for Spoon-billed Sandpipers declined steadily between 1950 and 2020: by 55.3% for total habitats, 53.7% for natural wetlands, and 58.2% for sandy beaches, most of which occurred between 1950 and 1980 (Fig. 3). Artificial wetlands expanded from 1950 and peaked around 1970, but 93.1% of that area had disappeared by 2020 (Fig. 3). Thus, although artificial wetlands may have temporarily supplemented the loss of natural wetlands between 1950 and 1980, both natural and artificial wetlands have declined continuously since 1980.

**Fig. 3.**
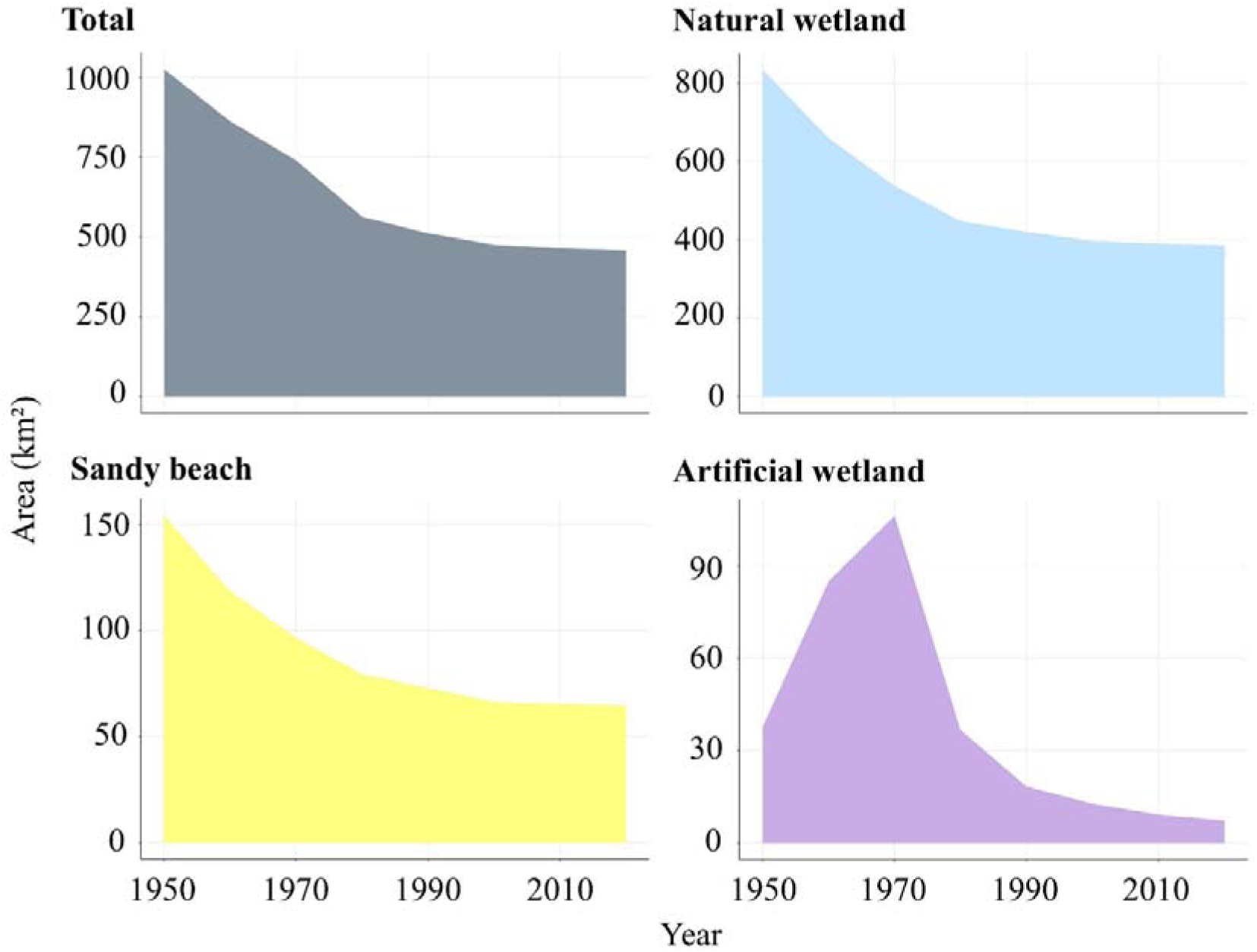
Temporal changes in area for three habitat types and the total across focal clusters from 1950 to 2020.

Consistent with patterns derived from observation records, sandpiper abundance per visit significantly decreased between the 1970s and 2010s across clusters, even after offsetting for cluster size and accounting for the significant positive associations with survey effort (Fig. 4; Table S7.1). In models using the opportunistic abundance as the response variable, abundance was significantly higher at higher latitudes, closer to breeding sites (Fig. 4; Table S7.1). Although both abundance models showed negative associations between abundance and habitat area across all three habitat types, abundance was positively associated with long-term cumulative change rates in natural wetlands and sandy beaches since 1950 (Fig. 4, 5AC; Table S7.1). Moreover, in both abundance models, abundance was lower at clusters with larger habitat extent that had experienced greater long-term habitat loss (Fig. 4, 5D; Table S7.1). Conversely, neither model showed a significant association with short-term change rates in total habitat area, sandy beaches, or artificial wetlands (Fig. 4, 5B; Table S7.1). These results suggest that spoonbill abundance was lower at clusters with greater long-term loss of natural habitats but was not related to short-term loss of natural and artificial habitats.

**Fig. 4.**
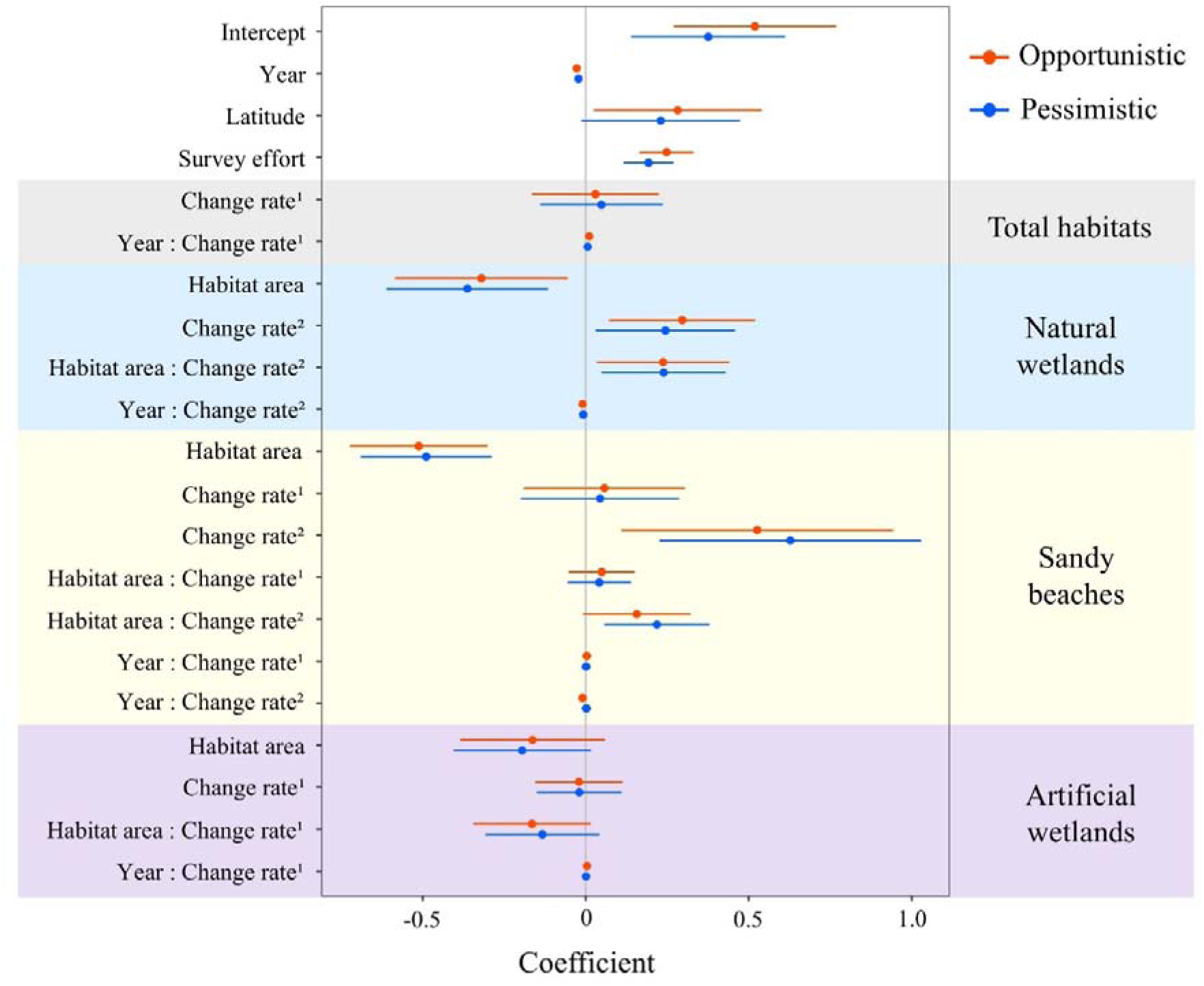
Effect sizes of variables from the two models using the core dataset. Change rate^1^: short-term change rate from one decade earlier; Change rate^2^: long-term change rate from 1950.

**Fig. 5.**
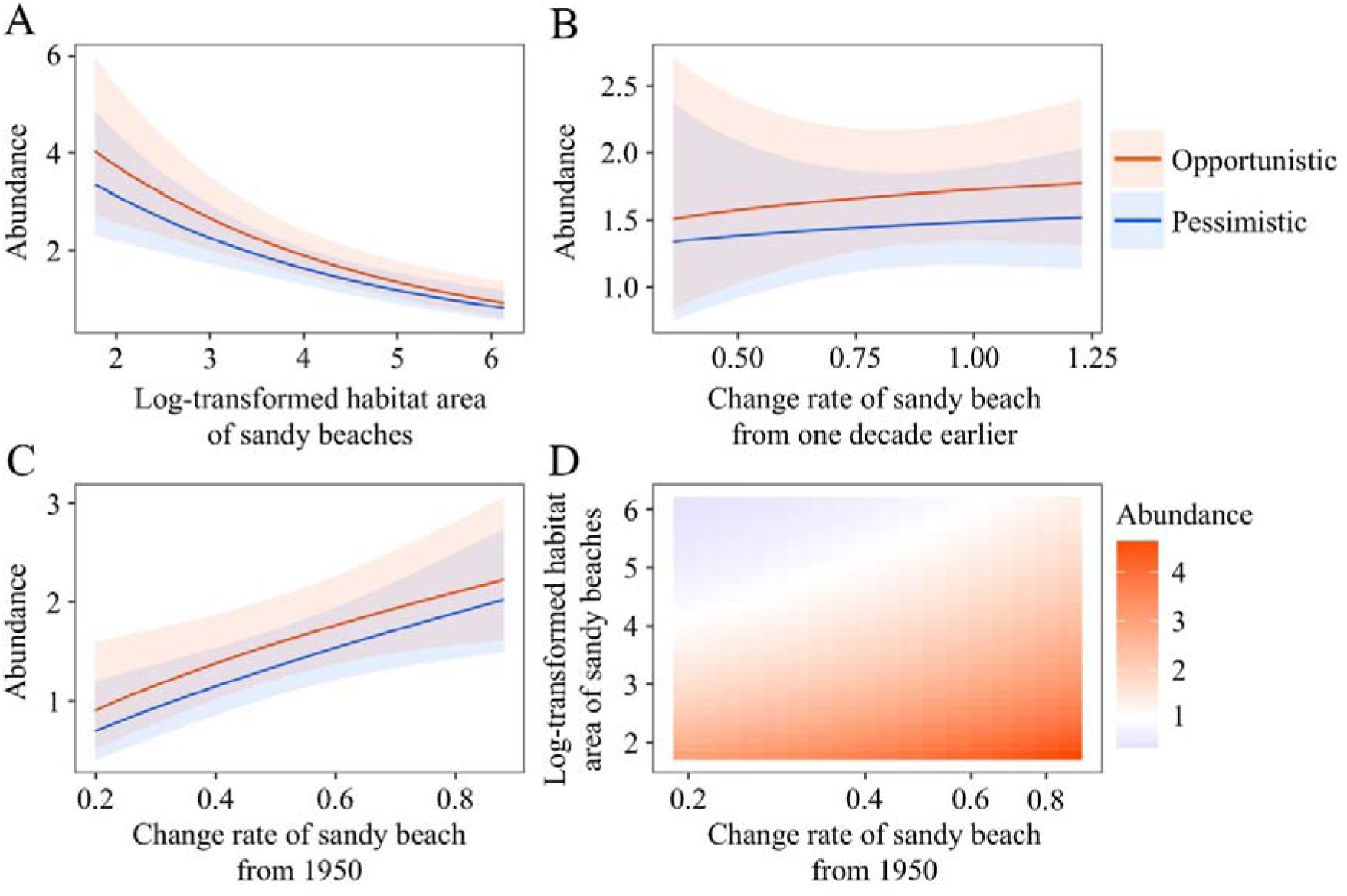
Response curves derived from models fitted to the core dataset. A) Habitat area of sandy beaches in observation; B) change rate of sandy beaches from one decade earlier; C) change rate of sandy beaches from 1950; and D) the interaction between habitat area of sandy beaches in observation and change rate of sandy beaches from 1950, derived from the model using opportunistic abundance.

## 4. Discussion

Stopover habitat quality and quantity are critical for the fitness and population persistence of long-distance migratory species (UNEP-WCMC, 2024), particularly those undergoing global declines or sensitive to environmental change (Habel et al., 2023). Yet few studies have simultaneously investigated how species abundance is associated with both short– and long-term spatiotemporal changes in natural and artificial stopover habitats. We examined the response of an endangered specialist migrant, Spoon-billed Sandpipers, to 70 years of natural and artificial habitat loss at stopover sites across Japan, while accounting for global year effects, uneven sampling effort, and regional variation in habitat size. Consistent with our hypothesis, historical loss of natural habitats was strongly associated with sandpiper abundance, with effect sizes varying across time and space.

Although our correlative analysis cannot establish causality between habitat loss and species populations, long-term changes in natural habitats were more strongly associated with sandpiper abundance than short-term changes. This highlights that focusing only on short-term habitat changes (e.g., within ∼ 10 years) may overlook or underestimate population responses; such patterns can be latent and emerge over longer time scales (Daskalova et al., 2020). Moreover, clusters with larger habitats that had experienced greater habitat loss tended to support fewer individuals, with abundance being lower rather than higher in larger habitats. The negative relationship between abundance and habitat size may indicate that Spoon-billed Sandpipers frequently select even small stopover habitats, with habitat selection potentially being more strongly influenced by migration routes, habitat quality, interspecific relationships, and weather conditions than by habitat size alone (Cohen et al., 2020; Flack et al., 2022; Schmaljohann et al., 2022). Furthermore, this pattern may also partly reflect limited data from larger wetlands, where Spoon-billed Sandpipers may occur within large mixed-species flocks, potentially reducing detection probability. Nevertheless, even if habitat extent is not a decisive factor for habitat selection at a given time, it may still be important over longer temporal scales. Accordingly, our findings provide empirical support for previous work indicating that loss of stopover habitats is likely to drive population declines in migratory species (Liu et al., 2022; Schmaljohann et al., 2022; Studds et al., 2017).

Spoon-billed Sandpipers also utilized artificial wetlands frequently; roughly one-sixth of observations occurred in artificial wetlands in our dataset, consistent with previous studies of other migratory shorebirds (Jackson et al., 2019, 2020). However, temporal changes in artificial wetlands were not associated with abundance. Two factors may explain this pattern. First, unlike natural wetlands and sandy beaches, the extent of artificial wetlands changed rapidly and inconsistently. Because short-lived wetlands may not sustain benthic communities and may provide less consistent prey availability, especially for specialists (Ma et al., 2004; Wang et al., 2022), our focal artificial wetlands may provide lower or less stable habitat quality than natural habitats. Second, the short-lived nature of artificial wetlands may weaken detectable relationships with abundance because populations may not increase in proportion to rapid habitat expansion. Although increasing artificial wetlands can be a cost-effective option for providing alternative habitats for generalist species where natural habitats are scarce (Aarif et al., 2025; Shimizu et al., 2025; Wang et al., 2022), protecting and enhancing natural wetland networks along flyways remains the more effective strategy for conserving endangered specialists such as Spoon-billed Sandpipers.

### 4.1. The impact of temporal changes in the quality and quantity of local stopover habitats on the global population of the species

Lower Spoon-billed Sandpiper abundance at stopover sites with greater natural habitat loss suggests these sites may have reduced stopover quality, function and/or attractiveness. Previous studies show that individuals utilizing poor-quality stopover habitats accumulate insufficient fat and lose body mass, leading to higher mortality (Baker et al., 2004; Rakhimberdiev et al., 2018). Although future work needs to clarify how quantitative and qualitative changes in local stopover habitats affect global population dynamics of the species, such local changes are likely to influence subsequent migration success, survival, and global population trends (Aarif et al. 2021; Schmaljohann et al., 2022; Studds et al., 2017).

Additionally, 89% of Spoon-billed Sandpipers whose age was assessed in our dataset were juveniles (Table S6.1). This high proportion may reflect Japan’s location at the eastern edge of the flyway. Juveniles are more likely than adults to use displaced routes, owing to inexperience and greater vulnerability to crosswinds (Flack et al., 2022; Sergio et al., 2014; Verhoeven et al., 2022). Thus, habitat loss at flyway edges may not affect all age classes equally but may disproportionately impact younger birds, contributing to their reduced survival and returning rates to breeding sites, as well as constraining population growth. Therefore, conserving peripheral stopover sites may be important for improving the migration success of young individuals and maintaining populations.

### 4.2. The broad-scale effects of habitat loss at stopover sites on migratory waterbirds

Sixty-two percent of our focal clusters overlap current shorebird monitoring sites, where large numbers of shorebirds—including nationally declining species (e.g., Red-necked stints) and globally declining species (e.g., Dunlins *C. alpina*)—occur during non-breeding and migration periods (MOEJ, 2023). This underscores that documented habitat loss at these sites could negatively affect habitat utilization by multiple shorebird species. However, interspecific differences in habitat preference, such as adaptability to artificial habitats, may produce different effect sizes of natural versus artificial wetland loss. For example, generalist and/or dominant species may be less negatively affected by natural habitat loss than specialists and may even increase in abundance due to competitive superiority within communities. Future research should account for interspecific differences in vulnerability to habitat loss and biotic interactions at stopover sites.

Concurrent historical habitat loss in other stopover regions is also likely to have carry-over effects on coastal shorebirds and to accelerate ongoing population declines (Figueiredo et al., 2019; Lisovski et al., 2024). For instance, during the same period of habitat loss in Japan, nearby stopover regions experienced comparable tidal wetland declines: 65.6% from the 1950s to 2000s in South Korea and 57.8% from the 1950s to 2010s in Taiwan (Chen et al., 2024; Murray et al., 2014). Additionally, declines in sandy beaches were associated with lower Spoon-billed Sandpiper abundance. Although previous studies have emphasized the conservation importance of tidal flats, especially for specific regions (Studds et al., 2017), protecting sandy beaches at stopover sites is also essential, particularly for Spoon-billed Sandpipers and other sympatric species.

### 4.3. Limitations

This research has two major limitations: one related to data sampling and the other related to the analytical framework. Regarding data sampling, since our analysis relied on opportunistic presence-only records, we could not fully account for zero truncation, unknown sampling efforts, and variation in detection probability, which would lead to underestimation of abundance and the impacts of habitat change. Nevertheless, our findings supported the hypothesis that stopover sites with greater habitat loss have lower abundance by incorporating survey effort as a covariate and implementing sensitivity analyses focused on data-rich periods, clusters, and seasons. In addition, our dataset appears to capture abundance and broad temporal trends reasonably. For example, our estimated global population size for the 1970s—derived from our dataset and population ratios of Japan to the world for the 1970s and 2020s—was 3755–10958, overlapping with 1578–8400 estimated in previous studies (Table S8.1).

A second analytical limitation is that our models used abundance as the response variable rather than population trends, which have been used in previous studies (Albaladejo_Robles et al., 2023; Spooner et al., 2018; Williams et al., 2022). As a result, our findings cannot be interpreted as evidence that stopover clusters experiencing greater habitat change also exhibited stronger population decreases (or increases). This limitation reflects the small and stochastic numbers of individuals observed in Japan, which made it difficult to estimate reliable population trends for each cluster. For example, Spooner et al. (2018) employed robust population trends fitted to generalized additive models or linear models based on criteria requiring data spanning more than four years and R^2^ ≥ 0.5; in our dataset, however, only a single cluster out of seventy met these criteria. Therefore, our approach may be particularly effective for specialist species that are rarely observed. By contrast, for more abundant species or well-sampled sites, population trends may be a more informative response variable.

## 5. Conclusion and conservation implications

Understanding how spatiotemporal changes in natural and artificial habitat extent at stopover sites affect populations is crucial for conserving migratory species, particularly specialists vulnerable to environmental change. Focusing on critically endangered Spoon-billed Sandpipers, we show that abundance was associated with greater cumulative natural habitat change, but not with short-term change and artificial habitat change. Our results suggest that abundance does not recover at the stopover sites that have experienced large-scale loss of natural habitats. Protecting remaining natural stopover habitats should be prioritized because their ecological functions cannot be offset by actions at breeding or non-breeding sites.

While artificial wetlands have recently gained attention as targets of habitat conservation (EAAFP, 2025), we should carefully examine whether their ecological functions are comparable to those of natural wetlands and improve the quality of artificial habitats to more closely approximate natural wetland functions. Although the temporary artificial wetlands examined in this study were not strongly associated with the abundance of our model species, larger and longer-lasting habitats can support higher biodiversity (Adler et al., 2005; Connor and McCoy, 1979), potentially enhancing their stopover functions. For instance, Japan is promoting the expansion of other effective area-based conservation measures (OECMs) to achieve the 30 by 30 target (UNEP-WCMC, 2025). Within this framework, managing coastal artificial wetlands created on reclaimed land for longer periods may be effective strategy. Nevertheless, future work should investigate why sensitive species use natural and artificial habitats differently and identify optimal spatial configurations and management for artificial wetlands.

## Data availability

Relating data and codes for analysis are available in https://doi.org/10.6084/m9.figshare.30007441.

## CRediT authorship contribution statement

**Takehiko Shimizu:** Conceptualization, Data curation, Formal analysis, Investigation, Methodology, Project administration, Software, Validation, Visualization, Writing – original draft. **Masayuki Senzaki:** Conceptualization, Data curation, Funding acquisition, Investigation, Supervision, Writing – original draft. **Munehiro Kitazawa:** Conceptualization, Writing – review and editing. **Minoru Kashiwagi:** Data curation, Writing – review and editing. **Hiroshi Tomida:** Data curation, Writing – review and editing.

## Acknowledgements

We used banding data with permission from the Yamashina Institute for Ornithology and Biodiversity Center of Japan. We thank regional birdwatchers for providing the observation records of Spoon-billed Sandpipers. We also appreciate the support in collecting observation records from Toshimitsu Nuka (Wild Bird Society of Japan) and Toshifumi Moriya (Bird Research).

## References

1. Aarif, K.M., Muzaffar, S.B., Babu, S., Prasadan, P.K., 2014. Shorebird assemblages respond to anthropogenic stress by altering habitat use in a wetland in India. Biodivers. Conserv. 23, 727–740. 10.1007/s10531-014-0630-9.

2. Aarif, K.M., Nefla, A., Nasser, M., Prasadan, P.K., Athira, T.R., Muzaffar, S.B., 2021. Multiple environmental factors and prey depletion determine declines in abundance and timing of departure in migratory shorebirds in the west coast of India. Glob. Ecol. Conserv. 26, e01518. 10.1016/j.gecco.2021.e01518.

3. Aarif, K.M., Nefla, A., Rubeena, K.A., Xu, Y., Musilova, Z., Musil, P., Wen, Lijia, Guo, Y., Naikoo, M.I., Sonne, C., Muzaffar, S.B., 2025. Rice fields as alternative foraging grounds: Rising shorebird diversity and abundance despite declines in natural coastal wintering sites. Ecol. Ind. 173, 113425. 10.1016/j.ecolind.2025.113425.

4. Adler, P.B., White, E.P., Lauenroth, W.K., Kaufman, D.M., Rassweiler, A., Rusak, J.A., 2005. Evidence for a general species-time-area relationship. Ecology. 86, 2032–2039. 10.1890/05-0067.

5. Albaladejo_Robles, G., Böhm, M., Newbold, T., 2023. Species life_history strategies affect population responses to temperature and land_cover changes. Glob. Chang. Biol. 29, 97–109. 10.1111/gcb.16454.

6. Aung, P. P., Buchanan, G.M., Round, P.D., Zöckler, C., Kelly, C., Tantipisanuh, N., & Gale, G.A., 2022. Foraging microhabitat selection of Spoon-billed Sandpiper in the Upper Gulf of Mottama, Myanmar. Glob. Ecol. Conserv. 35, e02077. 10.1016/j.gecco.2022.e02077.

7. Baker, A.J., Gonzalez, P.M., Piersma, T., Niles, L.J., Nascimento, I.L.S., Atkinson, P.W., Clark, N.A., Minton, C.D.T., Peck, M.K., & Aarts, G., 2004. Rapid population decline in red knots: fitness consequences of decreased refuelling rates and late arrival in Delaware Bay. Proc. Royal Soc. B: Biol. Sci. 271, 875–882. 10.1098/rspb.2003.2663.

8. BirdLife International, 2021. Calidris pygmaea. The IUCN Red List of Threatened Species 2021: e.T22693452A154738156. 10.2305/IUCN.UK.2021-3.RLTS.T22693452A154738156.en. (accessed 17 February 2025).

9. Bowler, D.E., Heldbjerg, H., Fox, A.D., O’hara, R.B., Böhning_Gaese, K., 2018. Disentangling the effects of multiple environmental drivers on population changes within communities. J. Anim. Ecol. 87, 1034–1045. 10.1111/1365-2656.12829.

10. Brooks, M.E., Kristensen, K., van Benthem, K.J., Magnusson, A., Berg, C.W., Nielsen, A., Skaug, H.J., Maechler, M., Bolker, M.M., 2017. glmmTMB balances speed and flexibility among packages for zero-inflated generalized linear mixed modeling. R. J. 9, 378–400. 10.32614/RJ-2017-066.

11. Chen, W.J., Chang, A.Y., Lin, C.C., Lin, R.S., Lin, D.L., Lee, P.F., 2024. Losing tidal flats at the midpoint of the East Asian-Australasian Flyway over the past 100 years. Wetlands 44, 59. 10.1007/s13157-024-01814-7.

12. Cheng, C., Ma, Z., 2023. Conservation interventions are required to improve bird breeding performance in artificial wetlands. Biol. Conserv. 278, 109872. 10.1016/j.biocon.2022.109872.

13. Choi, C., Gan, X., Hua, N., Wang, Y., Ma, Z., 2014. The habitat use and home range analysis of Dunlin (Calidris alpina) in Chongming Dongtan, China and their conservation implications. Wetlands 34, 255–266. 10.1007/s13157-013-0450-9.

14. Clark, N.A., Anderson, G.Q., Li, J., Syroechkovskiy, E.E., Tomkovich, P.S., Zöckler, C., Lee, R., & Green, R.E., 2018. First formal estimate of the world population of the Critically Endangered Spoon-billed Sandpiper Calidris pygmaea. Oryx 52, 137–146. 10.1017/S0030605316000806.

15. Cohen, E.B., Satterfield, D.A., 2020. ‘Chancing on a spectacle:’ co-occurring animal migrations and interspecific interactions. Ecography, 1657–1671. 10.1111/ecog.04958

16. Connor, E.F., McCoy, E.D., 1979. The statistics and biology of the species-area relationship. Am. Nat. 113, 791–833. 10.1086/283438.

17. Daskalova, G.N., Myers-Smith, I.H., Bjorkman, A.D., Blowes, S.A., Supp, S.R., Magurran, A.E., & Dornelas, M., 2020. Landscape-scale forest loss as a catalyst of population and biodiversity change. Science 368, 1341–1347. 10.1126/science.aba1289.

18. Dixon, J., 1918. The nesting grounds and nesting habits of the Spoon-billed Sandpiper. Auk 35, 387–404. 10.2307/4073213.

19. EAAFP, 2025. MOP12 Decision Papers DD.17 The EAAFP Initiative on Tidal Flats and Ecologically-associated Working Coastal Habitats (“EAAFP Tidal Flats Initiative – TFI”). https://eaaflyway.org/mop12-documents/ (accessed 14 January 2026).

20. Esri, 2024. ArcGIS Pro v2.4.0 (Version 2.4.0) [software]. https://www.esrij.com/products/arcgis-pro/.

21. Figueiredo, L., Krauss, J., Steffan-Dewenter, I., Cabral, J.S., 2019. Understanding extinction debt: spatio-temporal scales, mechanisms and a roadmap for future research. Ecography 42, 1973–1990. 10.1111/ecog.04740.

22. Flack, A., Aikens, E.O., Kölzsch, A., Nourani, E., Snell, K.R., Fiedler, W., Linek, N., Bauer, H.G., Thorup, K., Partecke, J., Wikelski, M., & Williams, H. J., 2022. New frontiers in bird migration research. Curr. Biol. 32, R1187–R1199. 10.1016/j.cub.2022.08.028.

23. GIAJ, 2023. Aerial Photographs Service. https://mapps.gsi.go.jp/maplibSearch.do#1. (assessed 15 August 2024)

24. Green, R.E., Syroechkovskiy, E.E., Anderson, G.Q.A., Chang, Q., Chowdhury, S., Clark, J.A., Foysal, M., Gerasimov, Y., Hughes, B., Kelly, C., Lappo, E., Lee, R., Leung, K.K.S., Li, J., Loktionov, E.Y., Melville, D.S., Phillips, J., Tomkovich, P.S., Weston, E., Weston, J., Yakushev, N., & Clark, N.A., 2021. New estimates of the size and trend of the world population of the Spoon-billed Sandpiper using three independent statistical models. Wader Study 128, 22–35. 10.18194/ws.00218.

25. Habel, J.C., Ulrich, W., Gros, P., Teucher, M., Schmitt, T., 2023. Butterfly species respond differently to climate warming and land use change in the northern Alps. Sci. Total Environ. 890, 164268. 10.1016/j.scitotenv.2023.164268.

26. Haddaway, N.R., Page, M.J., Pritchard, C.C., McGuinness, L.A., 2022. PRISMA2020: An R package and Shiny app for producing PRISMA 2020-compliant flow diagrams, with interactivity for optimised digital transparency and Open Synthesis. Campbell Syst. Rev. 18, e1230. 10.1002/cl2.1230.

27. Hale, R., Coleman, R., Pettigrove, V., Swearer, S.E., 2015. Identifying, preventing and mitigating ecological traps to improve the management of urban aquatic ecosystems. J. Appl. Ecol. 52, 928–939. 10.1111/1365-2664.12458.

28. Hale, R., Swearer, S.E., 2016. Ecological traps: current evidence and future directions. Proc. Royal Soc. B: Biol. Sci. 283, 20152647. 10.1098/rspb.2015.2647.

29. Hartig, F., 2022. DHARMa: Residual Diagnostics for Hierarchical (Multi-Level / Mixed) Regression Models. R package v0.4.6 (Version 0.4.6) [Software]. https://CRAN.R-project.org/package=DHARMa.

30. Horváth, Z., Ptacnik, R., Vad, C.F., Chase, J.M., 2019. Habitat loss over six decades accelerates regional and local biodiversity loss via changing landscape connectance. Ecol. Lett. 22, 1019–1027. 10.1111/ele.13260.

31. Ito, T., Hoshi, T., 2020. The Japanese Economy, second ed. MIT Press, Cambridge. https://mitpress.mit.edu/9780262538244/the-japanese-economy/.

32. Jackson, M.V., Carrasco, L.R., Choi, C.Y., Li, J., Ma, Z., Melville, D.S., Mu, T., Peng, H.B., Woodworth, B.K., Yang, Z., Zhang, L., & Fuller, R.A., 2019. Multiple habitat use by declining migratory birds necessitates joined-up conservation. Ecol. Evol. 9, 2505–2515. 10.1002/ece3.4895.

33. Jackson, M.V., Choi, C.Y., Amano, T., Estrella, S.M., Lei, W., Moores, N., Mundkur, T., Rogers, D.I., & Fuller, R.A., 2020. Navigating coasts of concrete: Pervasive use of artificial habitats by shorebirds in the Asia-Pacific. Biol. Conserv. 247, 108591. 10.1016/j.biocon.2020.108591.

34. Jaureguiberry, P., Titeux, N., Wiemers, M., Bowler, D.E., Coscieme, L., Golden, A.S., Guerra, C.A., Jacob, U., Takahashi, Y., Settele, J., Díaz, S., Molnár, Z., & Purvis, A., 2022. The direct drivers of recent global anthropogenic biodiversity loss. Sci. Adv. 8, eabm9982. 10.1126/sciadv.abm9982.

35. Jia, Y., Sun, L., Guo, J., Ren, S., Yang, H., Huang, G., Wen, L., Saintilan, N., Chen, Q., Wang, Y., & Lei, G., 2025. Identifying non-breeding habitat conservation gaps of the critically threatened Spoon-billed Sandpiper (*Calidris pygmaea*) using species distribution model. Glob. Ecol. Conserv. 61, e03640. 10.1016/j.gecco.2025.e03640.

36. Klaassen, R.H., Hake, M., Strandberg, R., Koks, B.J., Trierweiler, C., Exo, K.M., Bairlein, F., & Alerstam, T., 2014. When and where does mortality occur in migratory birds? Direct evidence from long_term satellite tracking of raptors. J. Anim. Ecol., 83, 176–184. 10.1111/1365-2656.12135.

37. Knapp, S., Schweiger, O., Kraberg, A., Asmus, H., Asmus, R., Brey, T., Frickenhaus, S., Gutt, J., Kühn, I., Liess, M., Musche, M., Pörtner, H., Seppelt, R., Klotz, S., & Krause, G., 2017. Do drivers of biodiversity change differ in importance across marine and terrestrial systems—Or is it just different research communities’ perspectives?. Sci. Total Environ. 574, 191–203. 10.1016/j.scitotenv.2016.09.002.

38. Lenth, R., 2024. emmeans: Estimated Marginal Means, aka Least-Squares Mean. R package v1.1.2 (Version 1.1.2) [Software]. https://CRAN.R-project.org/package=emmeans.

39. Le Provost, G., Badenhausser, I., Le Bagousse-Pinguet, Y., Clough, Y., Henckel, L., Violle, C., Bretagnolle, V., Roncoroni, M., Manning, P., & Gross, N., 2020. Land-use history impacts functional diversity across multiple trophic groups. Proc. Natl. Acad. Sci. 117, 1573–1579. 10.1073/pnas.1910023117.

40. Lisovski, S., Hoye, B.J., Conklin, J.R., Battley, P.F., Fuller, R.A., Gosbell, K.B., Klassen, M., Benjamin Lee, C., Murray, N.J., & Bauer, S., 2024. Predicting resilience of migratory birds to environmental change. Proc. Nat. Acad. Sci. 121, e2311146121. 10.1073/pnas.2311146121.

41. Liu, J., Lei, W., Mo, X., Hassell, C.J., Zhang, Z., Coulson, T., 2022. Unravelling the processes between phenotypic plasticity and population dynamics in migratory birds. J. Anim. Ecol. 91, 983–995. 10.1111/1365-2656.13686.

42. Lu, X., Yang, H., Piersma, T., Sun, L., Chen, Q., Jia, Y., Lei, G., Cheng, L., & Rao, X., 2022. Food resources for Spoon-billed Sandpipers (Calidris pygmaea) in the mudflats of Leizhou Bay, southern China. Front. Mar. Sci. 9, 1005327. 10.3389/fmars.2022.1005327.

43. Ma, Z., Li, B., Zhao, B., Jing, K., Tang, S., Chen, J., 2004. Are artificial wetlands good alternatives to natural wetlands for waterbirds? – A case study on Chongming Island, China. Biodivers. Conserv. 13, 333–350. 10.1023/B:BIOC.0000006502.96131.59.

44. MOEJ, 2023. Monitoring Site 1000. https://www.biodic.go.jp/moni1000/ (accessed 24 January 2023).

45. Montràs-Janer, T., Suggitt, A.J., Fox, R., Jönsson, M., Martay, B., Roy, D.B., Walker, K.J., & Auffret, A.G., 2024. Anthropogenic climate and land-use change drive short-and long-term biodiversity shifts across taxa. Nat. Ecol. Evol. 8, 739–751. 10.1038/s41559-024-02326-7.

46. Mu, T., Cai, S., Peng, H.B., Hassell, C.J., Boyle, A., Zhang, Z., Piersma, T., Wilcove, D.S., 2022. Evaluating staging habitat quality to advance the conservation of a declining migratory shorebird, Red Knot *Calidris canutus*. J. Appl. Ecol. 59, 2084–2093. 10.1111/1365-2664.14220.

47. Murray, N.J., Clemens, R.S., Phinn, S.R., Possingham, H.P., Fuller, R.A., 2014. Tracking the rapid loss of tidal wetlands in the Yellow Sea. Front. Ecol. Environ. 12, 267–272. 10.1890/130260.

48. Murray, N.J., Worthington, T.A., Bunting, P., Duce, S., Hagger, V., Lovelock, C.E., Lucas, R., Saunders, M., Sheaves, M., Spalding, M., Waltham, N.J., & Lyons, M. B., 2022. High-resolution mapping of losses and gains of Earth’s tidal wetlands. Science 376, 744–749. 10.1126/science.abm9583.

49. Newbold, T., Hudson, L.N., Hill, S.L.L., Contu, S., Lysenko, I., Senior, R.A., Börger, L., Bennett, D.J., Choimes, A., Collen, B., Day, J., Palma, A.D., Díaz, S., Echeverria-Londoño, S., Edgar, M.J., Feldman, A., Garon, M., Harrison, M.L.K., Alhusseini, T., Ingram, D.J., Itescu, Y., Kattge, J., Kemp, V., Kirkpatrick, L., Kleyer, M., Correia, D.L.P., Martin, C.D., Meiri, S., Novosolov, M., Pan, Y., Phillips, H.R.P., Purves, D.W., Robinson, A., Simpson, J., Tuck, S.L., Weiher, E., White, H.J., Ewers, R.M., Mace, G.M., Scharlemann, J.P.W., & Purvis, A., 2015. Global effects of land use on local terrestrial biodiversity. Nature 520, 45–50. 10.1038/nature14324.

50. Newbold, T., Hudson, L.N., Contu, S., Hill, S.L., Beck, J., Liu, Y., Meyer, C., Phillips, H.R.P., Scharlemann, J.P.W., & Purvis, A., 2018. Widespread winners and narrow-ranged losers: Land use homogenizes biodiversity in local assemblages worldwide. PLoS Biol., 16, e2006841. 10.1371/journal.pbio.2006841.

51. Northrup, J.M., Rivers, J.W., Yang, Z., Betts, M.G., 2019. Synergistic effects of climate and land_use change influence broad_scale avian population declines. Glob. Chang. Biol. 25, 1561–1575. 10.1111/gcb.14571.

52. O’Brien, R., 2007. A caution regarding rules of thumb for variance inflation factor. Qual. Quant. 41, 673–690. 10.1007/s11135-006-9018-6.

53. Page, M.J., McKenzie, J., Bossuyt, P.M., Boutron, I., Hoffmann, T.C., Mulrow, C.D., Shamseer, L., Tetzlaff, J.M., Akl, E.A., Brennan, S.E., Chou, R., Glanville, J., Grimshaw, J.M., Hróbjartsson, A., Lalu, M.M., Li, T., Loder, E.W., Mayo-Wilson, E., McDonald, S., McGuinness, L.A., Stewart, L.A., Thomas, J., Tricco, A., Welch, V.A., Whiting, P., & Moher, D., 2021. The PRISMA 2020 statement: an updated guideline for reporting systematic reviews. BMJ, 372. n71. 10.1136/bmj.n71.

54. Rajpar, M.N., Ahmad, S., Zakaria, M., Ahmad, A., Guo, X., Nabi, G., Wanghe, K., 2022. Artificial wetlands as alternative habitat for a wide range of waterbird species. Ecol. Indic. 138, 108855. 10.1016/j.ecolind.2022.108855.

55. Rakhimberdiev, E., Duijns, S., Karagicheva, J., Camphuysen, C.J., Dekinga, A., Dekker, R., Gavrilov, A., Horn, J.T., Jukema, J., Saveliev, A., Soloviev, M., Tibbitts, T.L., van Gils, J.A., & Piersma, T., 2018. Fuelling conditions at staging sites can mitigate Arctic warming effects in a migratory bird. Nat. Commun. 9, 4263. 10.1038/s41467-018-06673-5.

56. R Core Team, 2024. R: A Language and Environment for Statistical Computing v4.4.1 (Version 4.4.1). R Foundation for Statistical Computing, Vienna, Austria [Software]. https://www.R-project.org/.

57. Rushing, C.S., Ryder, T.B., Marra, P.P., 2016. Quantifying drivers of population dynamics for a migratory bird throughout the annual cycle. Proc. Royal Soc. B: Biol. Sci. 283, 20152846. 10.1098/rspb.2015.2846.

58. Schmaljohann, H., Eikenaar, C., Sapir, N., 2022. Understanding the ecological and evolutionary function of stopover in migrating birds. Biol. Rev. 97, 1231–1252. 10.1111/brv.12839.

59. Sergio, F., Tanferna, A., Stephanis, R.D., Jiménez, L.L., Blas, J., Tavecchia, G., Preatoni, D., Hiraldo, F. 2014. Individual improvements and selective mortality shape lifelong migratory performance. Nature 515, 410–413. 10.1038/nature13696.

60. Shimizu, T., Senzaki, M., Hori, S., Sueda, K., Ichihara, S., Ishida, R., Yoshigai, J., 2025. Short-term flooding in non-rice croplands provides stopover habitats for migrating waterbirds. Agric. Ecosyst. Environ. 383, 109504. 10.1016/j.agee.2025.109504.

61. Sievers, M., Hale, R., Parris, K.M., Swearer, S.E., 2018. Impacts of human_induced environmental change in wetlands on aquatic animals. Biol. Rev. 93, 529–554. 10.1111/brv.12358.

62. Spooner, F.E.B., Pearson, R.G., Freeman, R., 2018. Rapid warming is associated with population decline among terrestrial birds and mammals globally. Glob. Chang. Biol. 24, 4521–4531. 10.1111/gcb.14361.

63. Studds, C.E., Kendall, B.E., Murray, NJ., Wilson, H.B., Rogers, D.I., Clemens, R.S., Gosbell, K., Hassell, C.J., Jessop, R., Melville, D.S., Milton, D.A., Minton, D.A., Possingham, H.P., Riegen, A.C., Straw, P., Woehler, E.J., & Fuller, R.A., 2017. Rapid population decline in migratory shorebirds relying on Yellow Sea tidal mudflats as stopover sites. Nat. Commun. 8, 14895. 10.1038/ncomms14895.

64. Tomkovich, P.S., Syroechkovski Jr, E.E., Lappo, E.G., Zöckler, C., 2002. First indications of a sharp population decline in the globally threatened Spoon-billed Sandpiper Eurynorhynchus pygmeus. Bird Conserv. Int. 12, 1–18. 10.1017/S0959270902002010.

65. UNEP-WCMC., 2024. State of the World’s Migratory Species. UNEP-WCMC, Cambridge, United Kingdom. https://www.cms.int/en/publication/state-worlds-migratory-species-report.

66. UNEP-WCMC., 2025. Protected Area Profile for Japan from the World Database on Protected Areas. https://www.protectedplanet.net/country/JPN (accessed 19 December 2025).

67. Verhoeven, M.A., Loonstra, A.J., McBride, A.D., Kaspersma, W., Hooijmeijer, J.C., Both, C., Senner, N.R., & Piersma, T., 2022. Age_dependent timing and routes demonstrate developmental plasticity in a long_distance migratory bird. J. Anim. Ecol. 91, 566–579. 10.1111/1365-2656.13641.

68. Wang, X., Li, X., Ren, X., Jackson, M. V., Fuller, R. A., Melville, D. S., Amano, T., & Ma, Z., 2022. Effects of anthropogenic landscapes on population maintenance of waterbirds. Conserv. Biol. 36, e13808. 10.1111/cobi.13808.

69. Wild Bird Society of Japan Tokyo, 2003. The behavior of a tagged Spoon-billed Sandpiper arriving in the Tokyo Bay. Yurikamome, 566. 16–17. https://wbsjt.jimdoweb.com/.

70. Williams, J.J., Freeman, R., Spooner, F., Newbold, T., 2022. Vertebrate population trends are influenced by interactions between land use, climatic position, habitat loss and climate change. Glob. Chang. Biol. 28, 797–815. 10.1111/gcb.15978.

71. Zöckler, C., Syroechkovskiy, E.E., Atkinson, P.W., 2010. Rapid and continued population decline in Spoon-billed Sandpipers *Eurynorhynchus pygmeus* indicates imminent extinction unless conservation action is taken. Bird Conserv. Int. 20, 95–111. 10.1017/S0959270910000316.

72. Zöckler, C., Beresford, A.E., Bunting, G., Chowdhury, S.U., Clark, N.A., Fu, V.W.K., Htin Hla, T., Morozov, V.V., Syroechkovskiy, E., Kashiwagi, M., Lappo, E.G., Tong, M., Long, T.L., Yu, Y.T., Huettmann, F., Akasofu, H.K., Tomida, H., & Buchanan, G.M., 2016. The winter distribution of the Spoon-billed Sandpiper Calidris pygmaeus. Bird Conserv. Int. 26, 476–489. 10.1017/S0959270915000295.

73. Zurell, D., Graham, C.H., Gallien, L., Thuiller, W., Zimmermann, N.E., 2018. Long-distance migratory birds threatened by multiple independent risks from global change. Nat. Clim. Chang. 8, 992–996. 10.1038/s41558-018-0312-9.

74. Zuur, A.F., Ieno, E.N., Elphick, C.S., 2010. A protocol for data exploration to avoid common statistical problems: data exploration. Methods Ecol. Evol. 1, 3–14. 10.1111/j.2041-210X.2009.00001.x.

