## Supplementary material for "The impacts of seventy years of changes in stopover habitats on the critically endangered Spoon-billed Sandpiper *Calidris pygmaea*"

### **S1. Systematic collection of observation records of Spoon-billed Sandpiper**

**Table S1.1**. Overview and keywords used to collect observation records of Spoon-billed Sandpipers

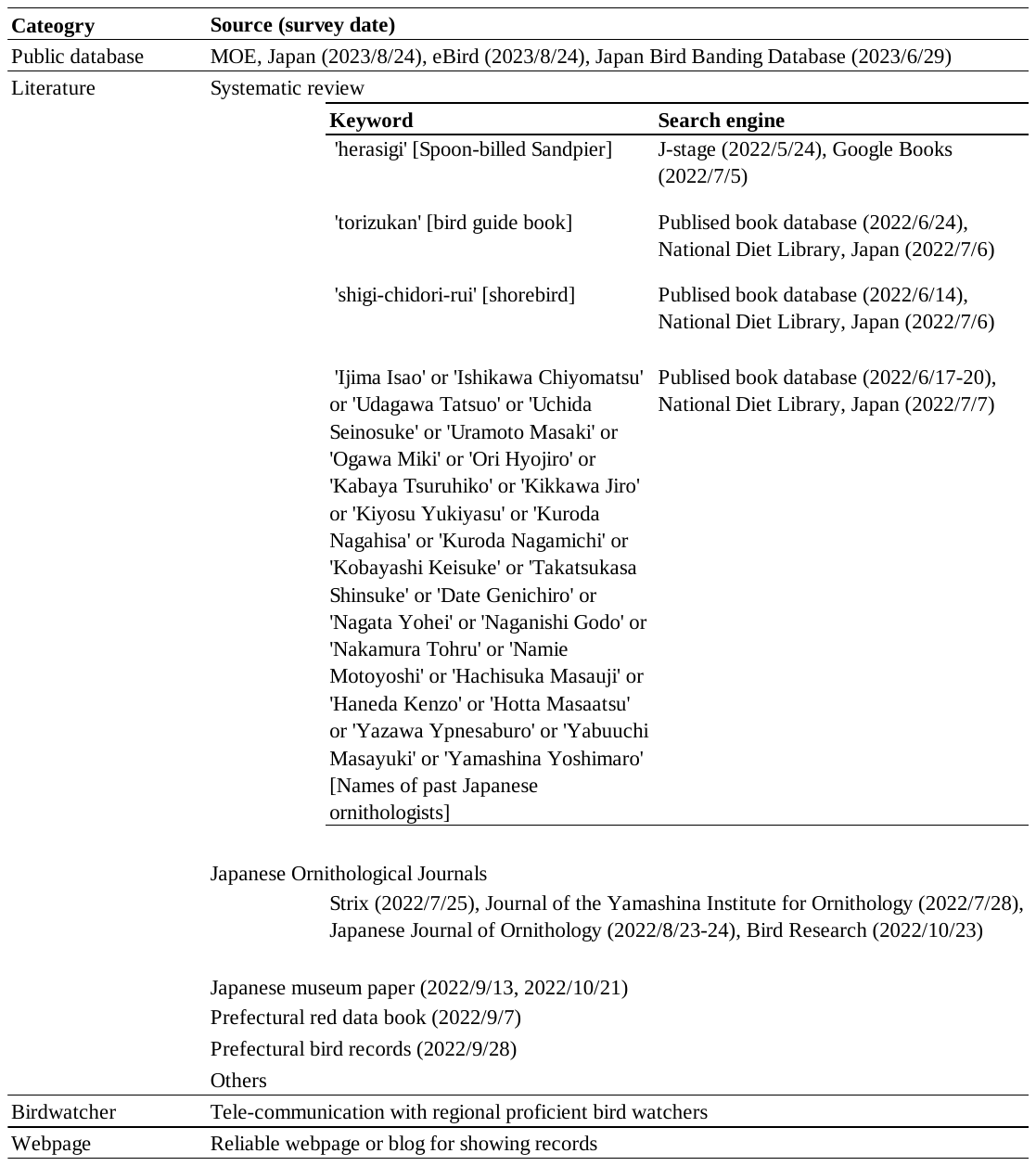

Dates in parentheses indicate when keyword searches were conducted.

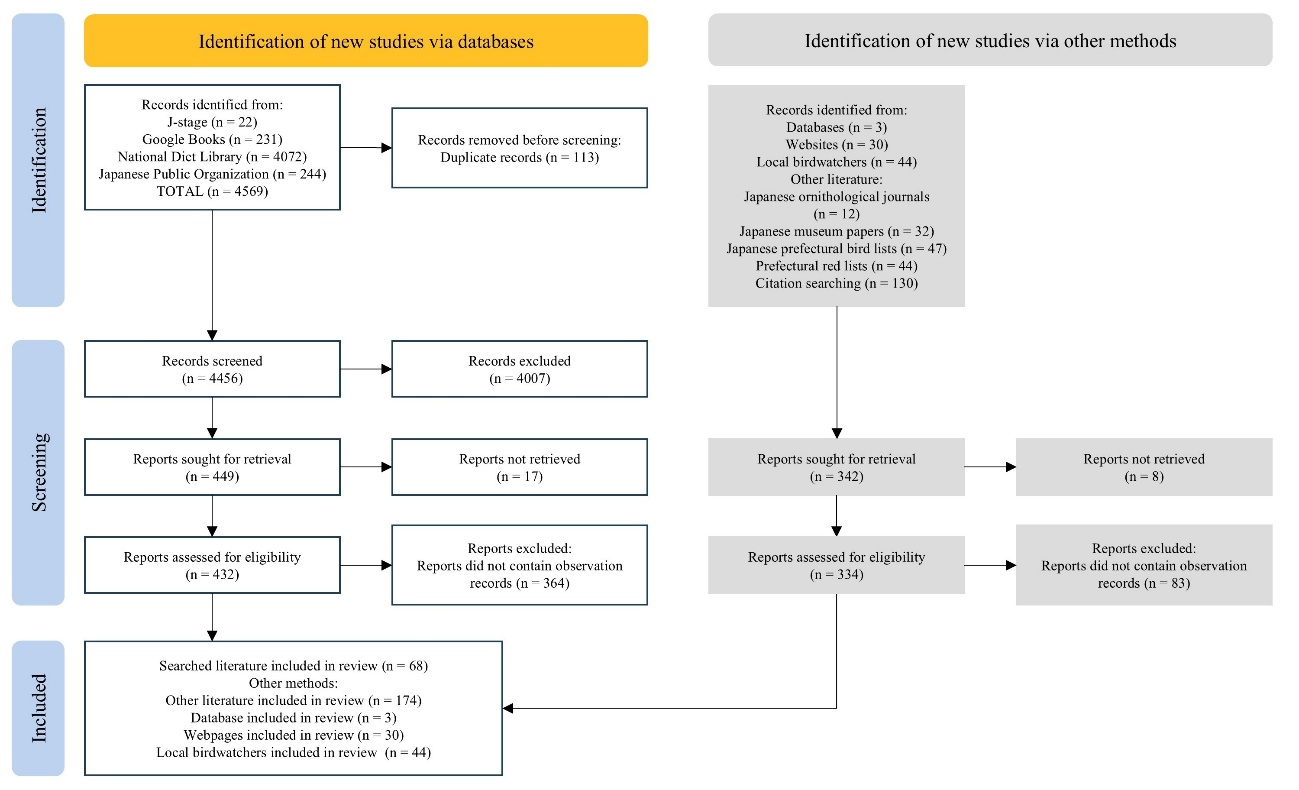

**Fig. S1.1.** PRISMA flow diagram for this study, adapted from Haddaway et al. (2022).

### **S2. Opportunistic and pessimistic abundance per stopover visit**

We considered records to be obvious duplicates when the following properties matched: prefecture, municipality, site name, year, month, day, and count. When more than one of these properties was missing, we classified the record as a potential duplicate (Table S2.1). To identify identical individuals, we considered the species’ movement range and stopover duration, although these are not well understood and may vary among individuals and sites. We relied on a study that tracked a single individual in Tokyo Bay (Wild Bird Society of Japan Tokyo, 2003). This study showed that the individual moved among multiple stopover sites up to about 30 km apart and remained in the area for about one month. Other records also indicate that individuals often remain at the same stopover site for several days to weeks (sometimes up to one month) and only rarely oversummer or overwinter. Based on these observations, we considered individuals to be identical when both criteria were met: 1) observations were made at sites < 30 km apart and 2) observation dates were < 15 days apart (Wild Bird Society of Japan Tokyo, 2003). When observation dates were ≥ 15 days but < 32 days apart, we classified individuals as potentially identical (Table S2.2). Complete criteria for identifying potentially identical individuals are provided in the Table S2.2. When source materials clearly reported movement ranges or lengths of stay that differed from these criteria, we followed the information provided in these materials. When aggregating counts for obviously or potentially identical individuals, we used the maximum number observed among the compared records. For records lacking an individual count, we assigned a value of one because at least one bird was confirmed present.

**Table S2.1**. Criteria for determining whether two or more records are different or obvious/potential duplicates.

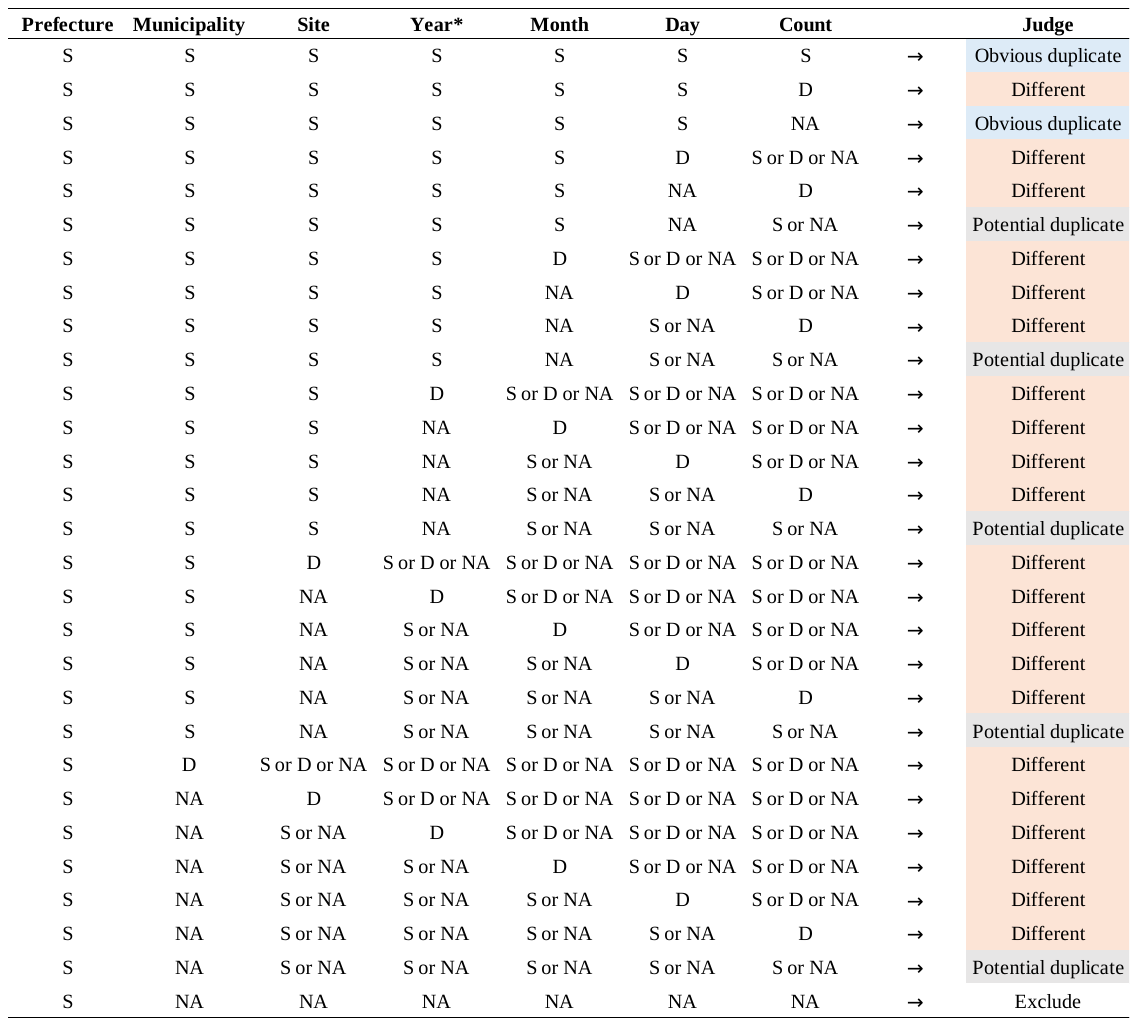

“S” = Same; “D” = Different; “NA” = missing information

**Table S2.2**. Criteria for determining whether two or more records represent different individuals or obvious/potentially identical individuals.

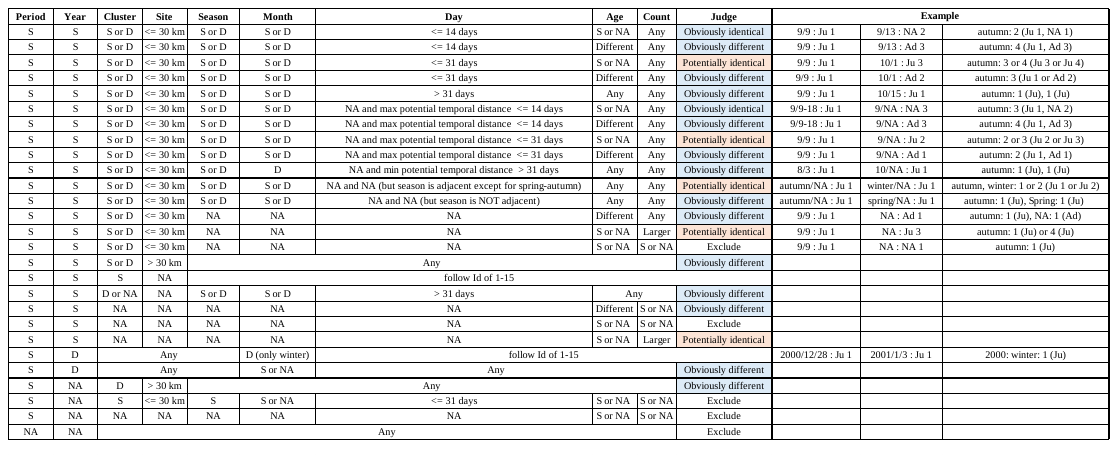
“S” = Same; “D” = Different; “NA” = missing information. “<= 30 km” = sites ≤30 km apart; “> 30 km” = sites >30 km apart; “<= 14 days” = date difference ≤14 days; “<= 31 days” = date difference >14 days and ≤31 days; “> 31 days" = date difference >31 days.

### **S3. Habitat types and habitat use by Spoon-billed Sandpipers**

**Table S3.1**. Number and proportion of records by habitat type used.

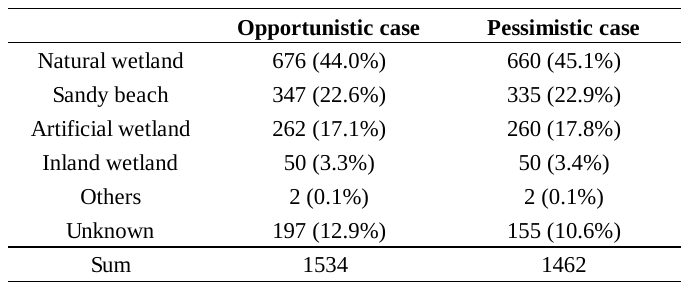

“Opportunistic” includes potential duplicate records; “pessimistic” excludes potential duplicates.

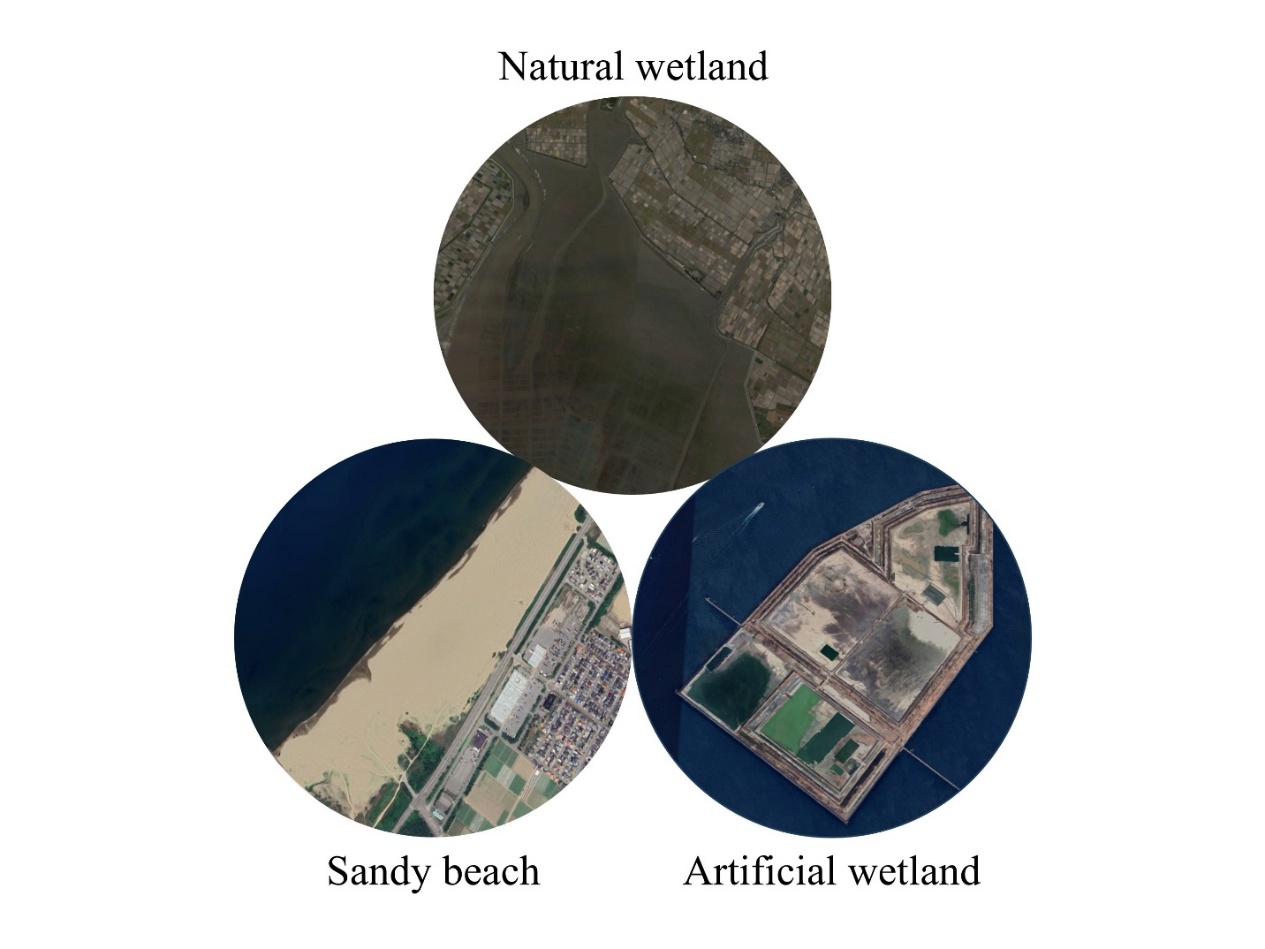

**Fig. S3.1.** The three focal habitat types. Images were generated in Google Earth Pro (https://support.google.com/earth/answer/21955?hl=en).

### **S4. Relationship between year and the global population of Spoon-billed Sandpipers**

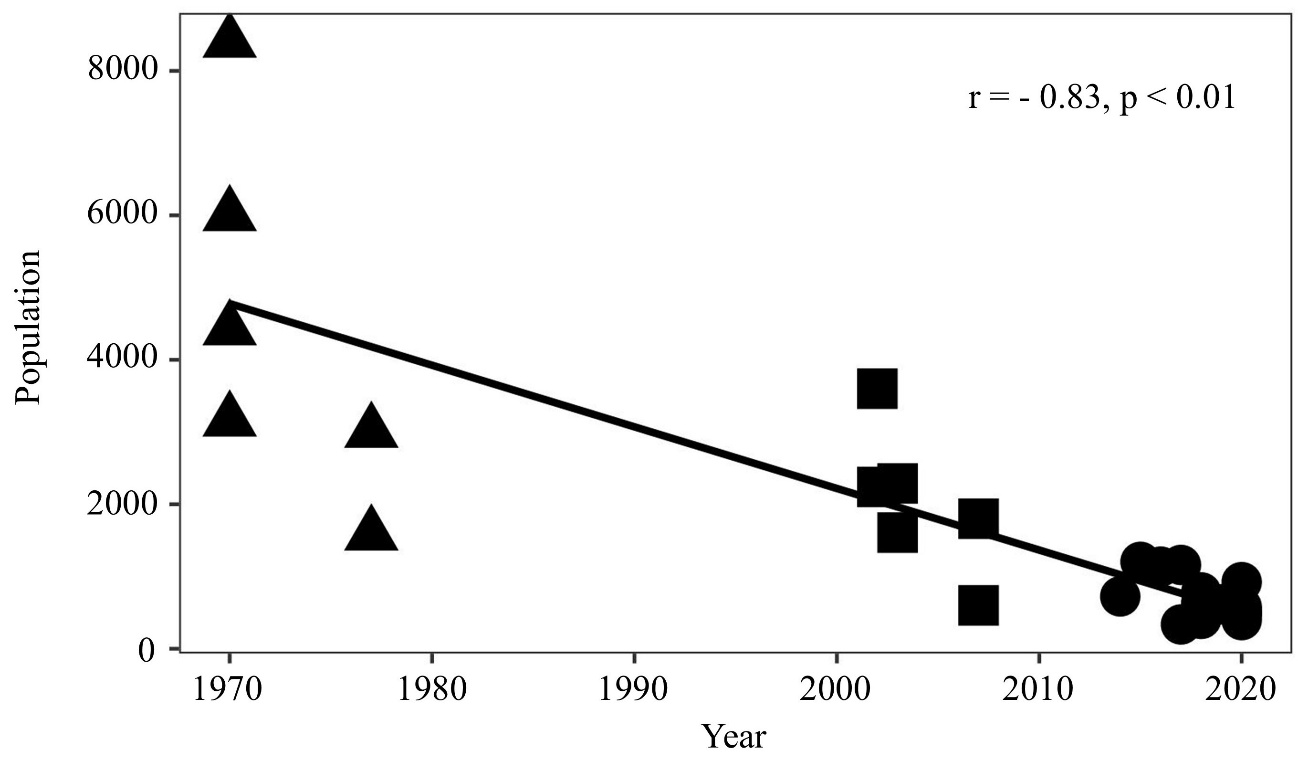

**Fig. S4.1.** Global population trend of the Spoon-billed Sandpiper. Circles: Green et al. (2021); squares: Zöckler et al. (2010); triangles: Tomkovich et al. (2002).

### **S5. Creating subset datasets for sensitivity analysis**

To test the sensitivity of our results, we compared outputs from multiple models using datasets that differed in four factors: cluster type, season, temporal window, and abundance definition (opportunistic/pessimistic). For cluster type, some clusters contained shorebird monitoring sites where regular surveys have been conducted annually in spring, autumn, and winter (MOEJ, 2023), increasing the likelihood that Spoon-billed Sandpipers were surveyed each year. We therefore created two datasets: one including all clusters and another including only monitoring clusters. For season, because most Spoon-billed Sandpipers were observed in autumn (approximately 85–86% of all individuals) (Table S5.1), we split the data into a dataset including all seasons and a dataset including only autumn records. For the temporal window, because the number of observation records was relatively stable between the 1970s and 2010s (Fig. S6.2A), we created two datasets: one spanning the 1960s-2020s and another spanning the 1970s-2010s. We used both opportunistic and pessimistic abundance accounting for potentially identical individuals. Combining these four factors, we constructed 16 abundance models (Table S5.2). We considered the two models using opportunistic and pessimistic abundance responses and the dataset restricted to monitoring clusters, autumn, and the 1970s-2010s to be the most robust.

**Table S5.1**. Number and proportion of observed individuals by season.

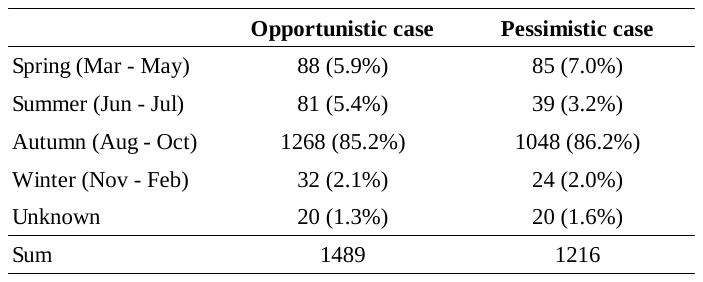

“Opportunistic” includes potential identical individuals; “pessimistic” excludes potential identical individuals.

**Table S5.2**. Setting for GLMMs

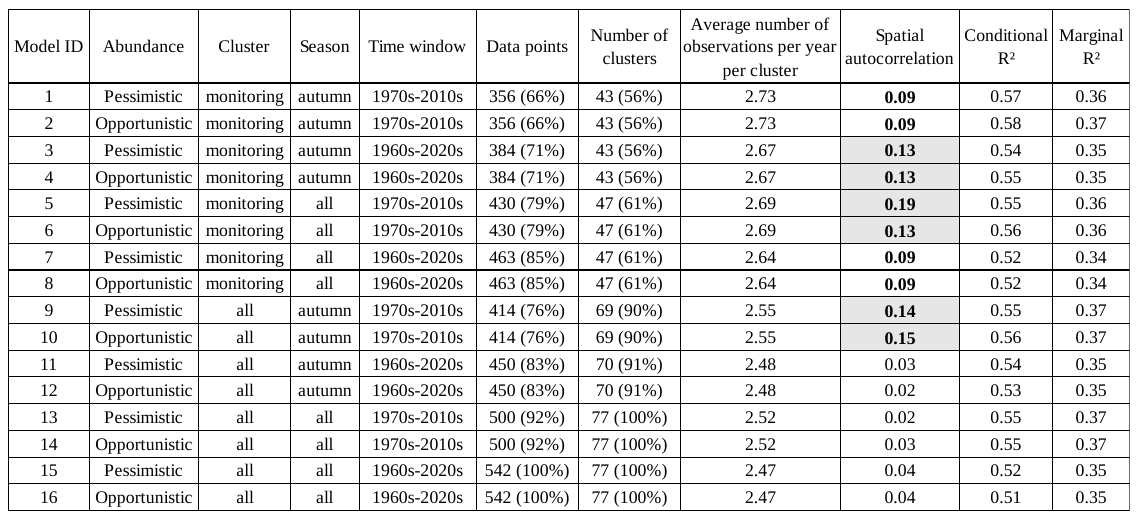

Model 1 and 2 utilized the core datasets, whereas the remaining models used subset datasets. “Opportunistic” includes potential identical individuals; “pessimistic” excludes potential identical individuals. For spatial autocorrelation, bold numbers indicate *p* < 0.05; gray cells indicate *p* < 0.01. The percentages of Data points and Number of clusters represent the ratios of values utilized in each model relative to the models using unfiltered datasets (i.e., model 15 and 16).

### **S6. Results of descriptive analyses**

**Table S6.1**. Age composition.

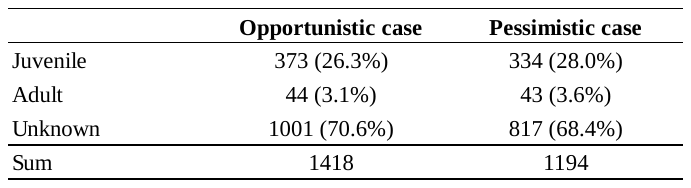

“Opportunistic” includes potential identical individuals; “pessimistic” excludes potential identical individuals.

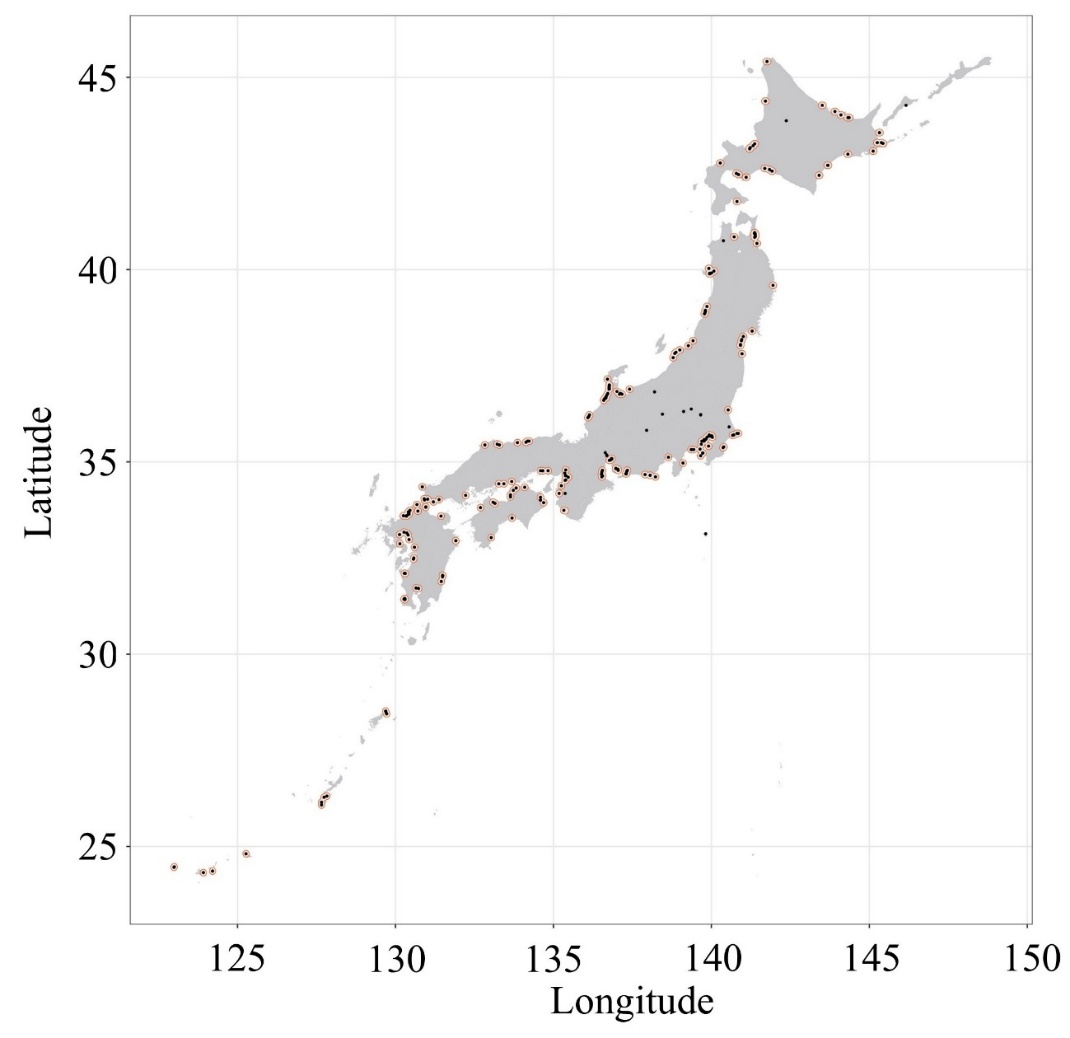

**Fig. S6.1.** Observed locations (black dots) and clusters (orange polygons) used for the habitat analysis.

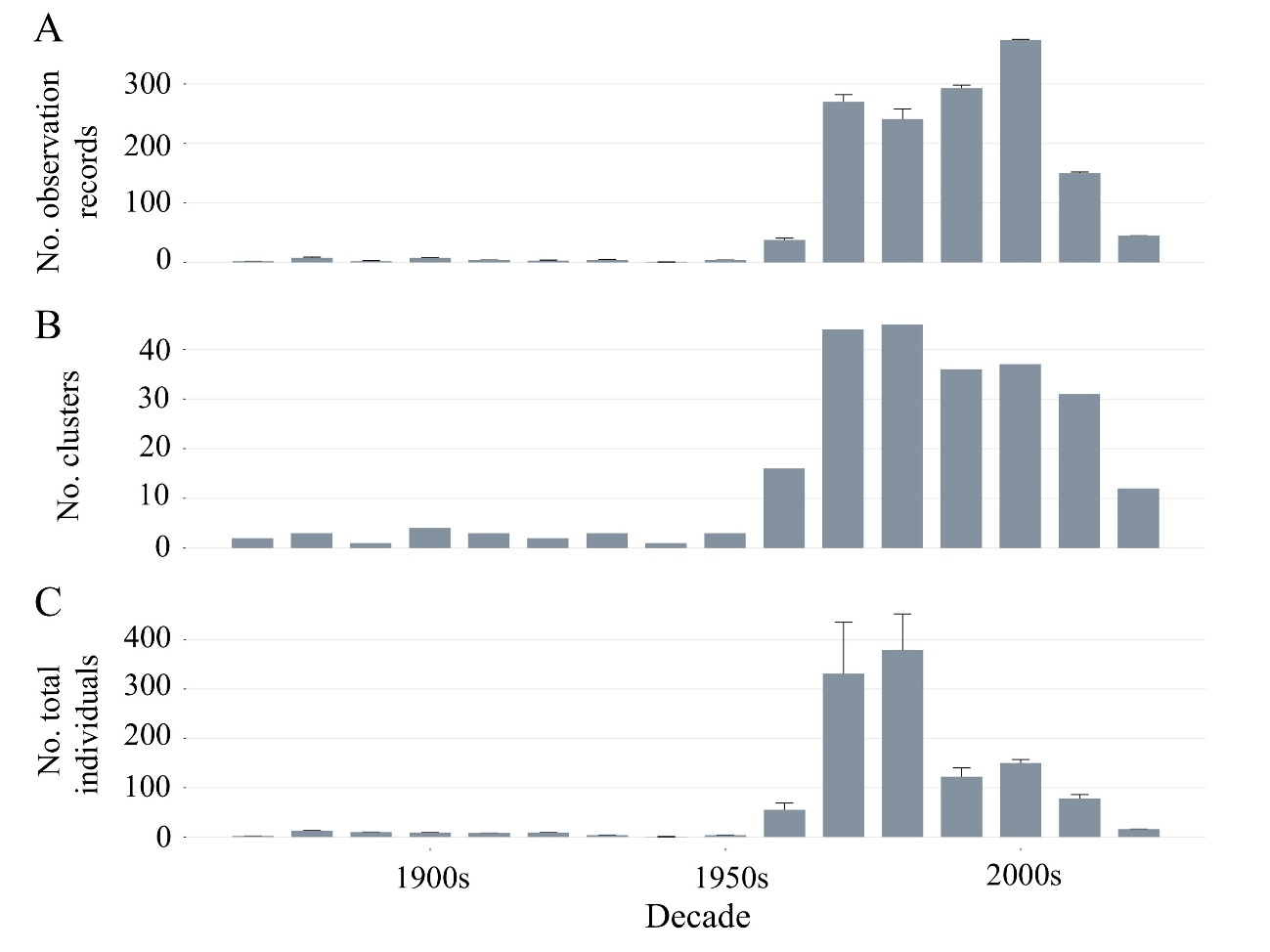

**Fig. S6.2.** Decadal trends in the number of observation records (A), observed clusters (B), and total individuals (C). Bar tops indicate the minimum values accounting for potential duplicates/identical individuals; error-bar tops indicate the maximum values without this consideration.

### **S7. Results of sensitivity analyses and GLMM coefficient tables**

The GLMMs used for sensitivity analysis showed consistent coefficient directions for explanatory variables, with only minor differences in effect sizes and *p*-values (Table S7.1). The following variables­­­—year, survey effort, habitat area of natural wetlands and sandy beaches—were significant (*p* < 0.05) in all models (16/16 models) (Table S7.1). Change rates for natural wetlands and sandy beaches from 1950, and the habitat area × change rate interaction for natural wetlands—significant in the core-dataset models—were also supported by most models (13, 15, and 15 models, respectively) (Table S7.1). In contrast, the year × change rate interaction for natural wetlands was significant only in model 2; thus, we considered this result not robust (Table S7.1). Some other variables showed results that varied by abundance definition and cluster subset. For latitude and the habitat area × change rate interaction for sandy beaches, significance differed between models with opportunistic abundance (6/8 and 2/8 models, respectively) and pessimistic abundance (4/8 and 8/8 models, respectively) (Table S7.1). In addition, models restricted to clusters with monitoring sites versus models including all clusters produced different results for variables related to artificial wetlands. Habitat area and change rate of artificial wetlands were not significant in the core-dataset models (model 1 and 2) but were significant in more models including all clusters (8/8 and 2/8 models, respectively) (Table S7.1). This may reflect that clusters with monitoring sites were less influenced by reclamation and therefore provided limited variation in artificial wetlands for explaining Spoon-billed Sandpiper abundance. Other variables not mentioned above were not consistently significant across models (Table S7.1).

**Table S7.1**. Results of GLMMs.

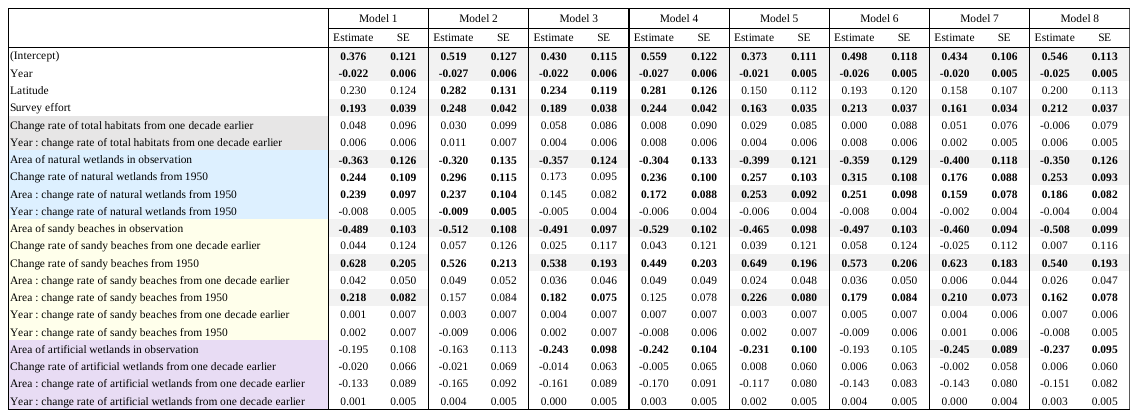

Model 1 and 2 utilized the core datasets, whereas the remaining models used subset datasets. Bold numbers indicate *p* < 0.05; gray cells indicate *p* < 0.01. Variable colors: gray = total habitat, sky blue = natural wetlands, yellow = sandy beaches, purple = artificial wetlands.

**Table S7.1**. Results of GLMMs (continued).

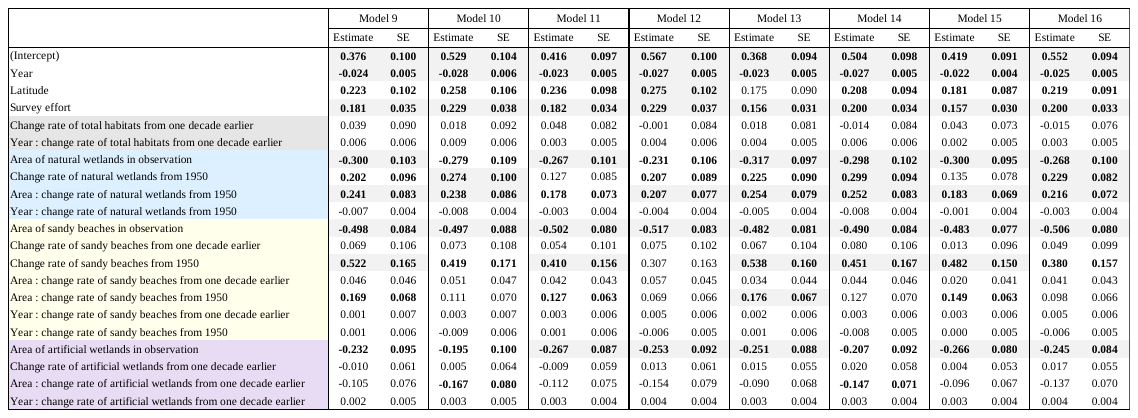

Model 1 and 2 utilized the core datasets, whereas the remaining models used subset datasets. Bold numbers indicate *p* < 0.05; gray cells indicate *p* < 0.01. Variable colors: gray = total habitat, sky blue = natural wetlands, yellow = sandy beaches, purple = artificial wetlands.

### **S8. Estimating the population size in the 1970s**

**Table S8.1**. Comparison of Spoon-billed Sandpiper population sizes in Japan and world (1970s vs. 2020s).

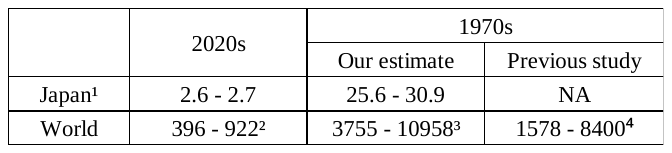

^1^ Japan’s population size was estimated as: marginal mean abundance per visit per site × average number of observed sites × average number of stopover visits per site, for the 1970s and 2020s, using the abundance models for all seasons and all clusters (i.e., model 1 and 9).

^2^ Global population in the 2020s from Green et al. (2021).

^3^ Global population in the 1970s was inferred from the Japan: world ratio using our 1970s/2020s estimates as follows:

$$N_{world}^{1970s} = {N_{Japan}^{1970s}\times N_{world}^{2020s}}/{N_{japan}^{2020s}}$$

where,

$N$: population size for the regions (world or Japan) in the periods (1970s or 2020s).

^4^ Because previous 1970s estimates ranged 6000–8400 (Tomkovich et al., 2002), we adopted 8400 as the maximum. Tomkovich et al. (2002) also indicated that ~1000 breeding pairs were present in the 1970s. Total population can be derived from breeding pairs or mature individuals using appropriate weights: Tomkovich et al. (2002) used ×3 (based on clutch size), whereas other studies incorporated juvenile survival (Clark et al. 2018; Green et al. 2021). Following Green et al. (2021), we used a weight of 1.578 (ratio of mature to total), yielding a minimum population estimate of $1000\times1.578=1578$ individuals.

### **References**

Clark, N.A., Anderson, G.Q., Li, J., Syroechkovskiy, E.E., Tomkovich, P.S., Zöckler, C., Lee, R., & Green, R.E., 2018. First formal estimate of the world population of the Critically Endangered spoon-billed sandpiper Calidris pygmaea. Oryx 52, 137–146. https://doi.org/10.1017/S0030605316000806.

Green, R.E., Syroechkovskiy, E.E., Anderson, G.Q.A., Chang, Q., Chowdhury, S., Clark, J.A., Foysal, M., Gerasimov, Y., Hughes, B., Kelly, C., Lappo, E., Lee, R., Leung, K.K.S., Li, J., Loktionov, E.Y., Melville, D.S., Phillips, J., Tomkovich, P.S., Weston, E., Weston, J., Yakushev, N., & Clark, N.A., 2021. New estimates of the size and trend of the world population of the Spoon-billed Sandpiper using three independent statistical models. Wader Study 128, 22–35. <https://doi.org/10.18194/ws.00218>.

Haddaway, N.R., Page, M.J., Pritchard, C.C., McGuinness, L.A., 2022. PRISMA2020: An R package and Shiny app for producing PRISMA 2020-compliant flow diagrams, with interactivity for optimised digital transparency and Open Synthesis. Campbell Syst. Rev. 18, e1230. https://doi.org/10.1002/cl2.1230.

Tomkovich, P.S., Syroechkovski Jr, E.E., Lappo, E.G., Zöckler, C., 2002. First indications of a sharp population decline in the globally threatened Spoon-billed Sandpiper Eurynorhynchus pygmeus. Bird Conserv. Int. 12, 1–18. <https://doi.org/10.1017/S0959270902002010>.

Wild Bird Society of Japan Tokyo, 2003. The behavior of a tagged Spoon-billed Sandpiper arriving in the Tokyo Bay. Yurikamome, 566. 16–17. https://wbsjt.jimdoweb.com/.

Zöckler, C., Syroechkovskiy, E.E., Atkinson, P.W., 2010. Rapid and continued population decline in the Spoon-billed Sandpiper Eurynorhynchus pygmeus indicates imminent extinction unless conservation action is taken. Bird Conserv. Int. 20, 95–111. https://doi.org/10.1017/S0959270910000316.
